# Glutamatergic neurons in the preoptic hypothalamus promote wakefulness, destabilize NREM sleep, suppress REM sleep, and regulate cortical dynamics

**DOI:** 10.1101/2020.10.20.347260

**Authors:** Alejandra Mondino, Viviane Hambrecht-Wiedbusch, Duan Li, A. Kane York, Dinesh Pal, Joaquin González, Pablo Torterolo, George A. Mashour, Giancarlo Vanini

## Abstract

Clinical and experimental data from the last nine decades indicate that the preoptic area of the hypothalamus is a critical node in a brain network that controls sleep onset and homeostasis. By contrast, we recently reported that a group of glutamatergic neurons in the lateral and medial preoptic area increases wakefulness, challenging the long-standing notion in sleep neurobiology that the preoptic area is exclusively somnogenic. However, the precise role of these subcortical neurons in the control of behavioral state transitions and cortical dynamics remains unknown. Therefore, in this study we used conditional expression of excitatory hM3Dq receptors in these preoptic glutamatergic (Vglut2+) neurons and show that their activation initiates wakefulness, decreases non-rapid eye movement (NREM) sleep, and causes a persistent suppression of rapid eye movement (REM) sleep. Activation of preoptic glutamatergic neurons also causes a high degree of NREM sleep fragmentation, promotes state instability with frequent arousals from sleep, and shifts cortical dynamics (including oscillations, connectivity, and complexity) to a more wake-like state. We conclude that a subset of preoptic glutamatergic neurons may initiate -but not maintain- arousals from sleep, and their inactivation may be required for NREM stability and REM sleep generation. Further, these data provide novel empirical evidence supporting the conclusion that the preoptic area causally contributes to the regulation of both sleep and wakefulness.

## Introduction

Since the beginning of the last century, the preoptic area of the hypothalamus has been considered a critical site for sleep generation. In the early 1900s, von Economo’s work revealed that patients suffering severe insomnia after encephalitis lethargica had extensive lesions within the rostral hypothalamus (von Economo, 1930). Consistent with this observation, subsequent studies demonstrated that lesioning of the preoptic area causes a prolonged, severe insomnia in cats (Sallanon et al., 1989) and rats (Nauta, 1946; John et al., 1994; Lu et al., 2000; Eikermann et al., 2011), and transplantation of fetal preoptic cells into the preoptic area partially restored sleep quantity in previously lesioned insomniac rats (John et al., 1998). Infusion of adenosine (or adenosine analogs) into the preoptic region increases sleep and decreases wakefulness (Ticho and Radulovacki, 1991; Mendelson, 2000), whereas pharmacologic inhibition of this region reduces sleep (Alam and Mallick, 1990; Lin et al., 1989; Benedetto et al., 2012). Additionally, evidence from single-cell recording and cFos studies confirmed that the median preoptic nucleus (MnPO) as well as ventrolateral and medial preoptic area contain neurons that are mainly active during non-rapid eye movement (NREM) sleep (Koyama and Hayaishi, 1994; Szymusiak et al., 1998; Suntsova et al., 2002; Takahashi et al., 2009; Sakai, 2011; Alam et al., 2014; Zhang et al., 2015; Chung et al., 2017; Harding et al., 2018) and, to a lesser extent, during rapid eye movement (REM) sleep (Koyama and Hayaishi, 1994; Lu et al., 2002; Suntsova et al., 2002; Gvilia et al., 2006; Dentico et al., 2009; Sakai, 2011; Alam et al., 2014). Importantly, a subset of these cells increases its activity during periods of prolonged wakefulness with increased sleep pressure, suggesting a role in the regulation of both NREM and REM sleep homeostasis (Gvilia et al., 2006; Dentico et al., 2009; Todd et al., 2010; Alam et al., 2014; Gvilia et al., 2017).

GABAergic neurons in the median, ventrolateral, and lateral-medial preoptic area are sleep-active (most co-express galanin) and innervate monoaminergic arousal-promoting systems (Sherin et al., 1998; Gaus et al., 2002; Uschakov et al., 2006; Hsieh et al., 2011; Chung et al., 2017). Activation of preoptic GABAergic and galaninergic neurons promotes NREM sleep (Chung et al., 2017; Harding et al., 2018; Kroeger et al., 2018; Vanini et al., 2020), and lesion (Lu et al., 2000; Ma et al., 2019) or neuromodulation of these neurons (Harding et al., 2018; Kroeger et al., 2018) alters the electroencephalogram (EEG) during sleep-wake states. Furthermore, a recent study using activity-dependent tagging and subsequent stimulation of previously tagged neurons showed that a subgroup of glutamatergic neurons within the median and medial preoptic nucleus of the hypothalamus promotes body cooling and NREM sleep (Harding et al., 2018). However, contrary to the prevailing idea in the field that virtually all preoptic neurons that regulate sleep-wake states are somnogenic, we recently reported that a region including the ventrolateral preoptic nucleus (VLPO) and the ventral half of the medial and lateral preoptic area, collectively referred in the current study as medial-lateral preoptic region, contains glutamatergic neurons that promote wakefulness (Vanini et al., 2020).

Here we used a chemogenetic stimulation approach to investigate further the role of these preoptic glutamatergic neurons in the regulation of sleep-wake states, state transitions, and cortical dynamics. We show that stimulation of glutamatergic neurons in the medial-lateral preoptic region causes a transient increase in wakefulness, and a decrease in both NREM and REM sleep. Interestingly, activation of these neurons produces a high degree of NREM sleep fragmentation and state instability, a “lighter” NREM sleep state, and a long-lasting suppression of REM sleep. Thus, our data suggest that a subset of preoptic glutamatergic neurons may initiate -but not maintain- arousal from sleep, and their inactivation might be required for NREM stability and REM sleep generation. Furthermore, these results provide novel empirical evidence that the preoptic area plays a dual role in the regulation of both sleep and wakefulness.

## Methods

### Mice

Vglut2-IRES-Cre (Slc17a6tm2(cre)Lowl/J; Stock# 016963) mice were purchased from Jackson Laboratory, bred at the University of Michigan animal care facility, and genotyped (Transnetyx) before weaning. Adult male mice used in this study (n = 28) were housed in groups in a 12-h light:dark cycle (lights on at 6:00 AM) with free access to food and water. All experiments were approved by the Institutional Animal Care and Use Committee and the Institutional Biosafety Committee, and were conducted in accordance with the recommendations published in the Guide for the Care and Use of Laboratory Animals (Ed 8, National Academies Press, Washington, D.C., 2001).

### Viral vector and drugs

For Cre-recombinase dependent expression of the excitatory designer receptor hM3Dq in preoptic glutamatergic (Vglut2+) neurons we used the adeno-associated viral vector AAV-hSyn-DIO-hM3D(Gq)-mCherry (Addgene; Cat.# 50459-AAV5) (Krashes et al., 2011; Vanini et al., 2020). In a separate group of mice used for control experiments, we used the Cre-dependent vector AAV-hSyn-DIO-mCherry (Addgene; Cat.# 50459-AAV5) that lacked the coding sequence for the designer receptor and only contained the fluorescent reporter mCherry. The titer of the viral solutions was 3.7 to 7.8 × 10^12^ copies per ml. Dimethyl sulfoxide (DMSO) and clozapine-N-oxide (CNO; agonist at hM3Dq receptors) were purchased from Sigma-Aldrich (St. Louis, MO, USA). A stock solution of CNO (0.1 mg/ml in saline solution containing 0.5% DMSO was prepared and stored in aliquots, and then frozen at−20 °C until use. Before each experiment, an aliquot of the stock solution was thawed at room temperature protected from the light until injection time. Mice received 1 mg/kg CNO or a vehicle control solution (saline with 0.5% DMSO) by intraperitoneal injection (i/p); the injection volume was 0.1 mL per 10 g of body weight (Vanini et al., 2020).

### Viral injection and electrode implantation for sleep studies

The procedure for vector injection into the preoptic area was performed as described previously (Vanini et al., 2020). Mice were anesthetized in an induction chamber with 5.0% isoflurane (Hospira, Inc., Lake Forest, IL, USA) in 100% O2. The delivered concentration of isoflurane was monitored continuously by spectrometry (Cardiocap/5; Datex-Ohmeda, Louisville, CO, USA). Following anesthetic induction, mice received pre-emptive analgesia (5 mg/kg carprofen; subcutaneous), and were immediately placed in a Kopf Model 962 stereotaxic frame fitted with a mouse adaptor and a mouse anesthesia mask (Models 922 and 907, respectively) (David Kopf Instruments, Tujunga, CA, USA). Thereafter, the concentration of isoflurane was reduced and maintained at 1.6 to 2.0% throughout the surgical procedure. Core body temperature was maintained at 37 – 38 °C with a water-filled pad connected to a heat pump (Gaymar Industries, Orchard Park, NY, USA). For the medial-lateral preoptic region, vector injections were performed bilaterally (36 nL on each side); stereotaxic coordinates: AP = +0.15 mm, ML = ±0.5 mm and DV = −5.0 mm from bregma (Vanini et al., 2020). Injections were performed at a rate of 5 nL/min using a Hamilton Neuros Syringe 7000 (5 mL; Hamilton Company, Reno, NV, USA) mounted on a microinjection syringe pump, driven by a digital Micro2T controller (Model UMP3T-2; World Precision Instruments, Sarasota, FL, USA). After each injection, the syringe was maintained in position for an additional 5 minutes to avoid vector reflux. We previously confirmed that this volume, stereotaxic coordinates and injection procedure yielded a reliable expression of hM3Dq receptors in the target region (Vanini et al., 2020). The scalp incision was closed with sutures and mice were placed under a controlled heat source until full recovery. Mice were then returned to the vivarium and group-housed with their respective littermates. Analgesia was maintained with carprofen (5 mg/kg every 24 h) for a minimum of 48 hours after surgery.

Three weeks following vector injection, mice were anesthetized with isoflurane and EEG electrodes (8IE3632016XE, E363/20/1.6/SPC; Plastics One, Roanoke, VA, USA) were implanted above the right frontal (AP = +1.5 mm, ML = +2.0 mm from bregma) and occipital (AP = −3.2 mm, ML = +3.0 mm from bregma) cortices. A reference screw electrode was implanted over the cerebellum, and a pair of electromyogram (EMG) electrodes (8IE36376XXXE, E363/76/SPC; Plastics One) was inserted bilaterally into the dorsal neck muscles. All electrode pins were then inserted into an electrode pedestal (MS363; Plastics One) that was affixed to the skull with dental cement (Fast Cure Powder/Liquid, Product# 335201; GC America, Inc., Alsip, IL, USA). Thereafter, the delivery of isoflurane was discontinued, and mice were allowed to recover following the same postoperative care protocol outlined above. Mice were allowed a minimum of 10 days before experiments began.

### EEG/EMG data acquisition

At least 1 week after the surgery for implantation of EEG/EMG electrodes, mice were conditioned to handling, simulating drug injections and tethering in the recording setup for 5 days. On the day of the experiment, mice received an injection of vehicle or CNO and EEG/EMG signals were recorded for 6 hours (injection and recording start time: 10:00 AM). Monopolar EEG and bipolar EMG signals were amplified (X 1000) and digitized (sampling rate = 1024 Hz), respectively, with a Model 1700 AC amplifier (A-M Systems, Sequim, WA, USA) and a Micro3 1401 acquisition unit and Spike2 software (Cambridge Electronic Design, Cambridge, UK); notch filters were used for each mouse recording (both post-vehicle and CNO) when 60 Hz electrical noise was present. All signals were bandpass filtered between 0.1 – 500 Hz (EEG) and 10 – 500 Hz (EMG). Mouse behaviors were video recorded, in synchrony with the electrophysiologic recordings in Spike2.

### Analysis of sleep-wake states

States of wakefulness, NREM, and REM sleep were manually scored in Spike2 in 5-second epochs using standard criteria described in a previous publication (Vanini et al., 2020). Wakefulness was defined by low-amplitude, high-frequency EEG activity accompanied by an active EMG characterized by high tone with phasic movements. NREM sleep was recognized by high-amplitude and low-frequency EEG waveforms along with a low muscle tone. REM sleep was identified by low-amplitude, high-frequency EEG signals with prominent, regular theta rhythm (particularly evident in the occipital cortex), and EMG atonia. Total time spent in wakefulness, NREM and REM sleep, the average duration and number of bouts for each vigilance state, was analyzed over the 6-hour recording period, as well as in 1-hour bins to assess the mean duration of the treatment effect.The mean latency to NREM sleep and REM sleep onset was compared between treatment conditions. Additionally, we reanalyzed sleep recordings from a group of Vglut2-Cre mice used in a recent study with confirmed hM3Dq receptor expression within the MnPO of the hypothalamus (Vanini et al., 2020). The viral vector, surgical procedures, experimental design and dose of CNO were identical in both studies. These mice served in the current study as a site-control group.

### Sleep fragmentation analysis

An initial exploratory analysis revealed that chemogenetic activation of preoptic glutamatergic neurons substantially increased the number and decreased the duration of wakefulness and NREM sleep bouts, revealing a robust sleep fragmentation. Therefore, to further characterize this phenomenon we calculated the stability of wake and NREM sleep states, between- and within-state transition probabilities, as well as a fragmentation index. First, the mean number of transitions from NREM to wakefulness was quantified over the 6-hour recording period and for every 1-hour bin. The stability of each state was calculated using a Markov memoryless model. This is a regression-like method in which variables are modeled as a function of the previous observations (Kim et al., 2009; Perez-Atencio et al., 2018). In this formalism, the probability of being in one state is determined by the previous state and the transition probability from one state to another. The probability of each specific state was calculated by the number of times that this state was scored over the total number of epochs recorded and the transition probability was obtained as the conditional probability P(X/Y), i.e., the number of times that state Y transitioned to state X in the following epoch, divided by the number of epochs of the Y state. Next, we calculated a fragmentation index (FI) for wakefulness and NREM sleep (codes are available at https://github.com/joaqgonzar/Fragmentation_Index). We defined FI as FI = 1 - p(X/X), i.e., 1 minus the probability of transitioning from the state X to the same state X. This implies that FI = 1 only if the state is completely fragmented, in other words, if the probability of transitioning to the same state equals to 0. Finally, as the sleep-wake cycle is characterized by bouts of short and long duration, we generated histograms of bout length for wakefulness and NREM sleep. These plots were obtained by counting the number of events that occurred in each increment of 10 s for wakefulness and 50 s for NREM sleep.

### Determination of changes in body temperature after activation of glutamatergic neurons in the medial-lateral preoptic region

In a separate group of mice, core body temperature was measured using a lubricated mouse rectal probe (Model# 600-1000, Barnant Company, Barrington, IL, USA). Temperature measures were obtained prior to the injection of CNO (1.0 mg/kg) and in 10-min intervals for 90 min post-injection. If a temperature drop was detected, mice were placed under a heating lamp in their respective home cages between testing intervals.

### EEG signal processing and analysis

EEG analysis (power, coherence, connectivity, and complexity; 0.1 – 115 Hz) was conducted on artifact-free, non-transition epochs during the initial 3-hour block after vehicle and CNO injection. Digitized, raw EEG signals in frontal and occipital channels were exported from Spike2 software into MATLAB (version 2019b; MathWorks, Inc., Natick, MA) and downsampled to 512 Hz (resample.m function in Matlab signal processing toolbox). Because notch filters were applied to EEG channels in some recordings (always in vehicle/CNO pairs, for consistency), frequencies between 45 – 75 Hz were not considered for the analysis. The power-line interference (60 Hz and 120 Hz), if present, was removed using multi-taper regression technique and Thomas F-statistics implemented in CleanLine plugin for EEGLAB toolbox (Delorme and Makeig, 2004). The 3-hour long recording blocks were segmented into 5-s windows with no overlap. All EEG measures were calculated in each window and then averaged over all available windows to obtain the mean values for each vigilance state, treatment condition and mouse.

#### Spectral power and coherence analysis

The EEG power spectrum was estimated using the multitaper method in Chronux analysis toolbox (Mitra and Bokil, 2008) with a window length = 5 s without overlap, time-bandwidth product = 2, and number of tapers = 3. For each mouse in each treatment group, the mean power spectrum in each behavioral state (wake, NREM, REM) was obtained by averaging the power spectra across all available windows in frontal and occipital channels, respectively. Furthermore, the power values were calculated for slow oscillations (0.5 – 1 Hz) as well as delta (1 – 4 Hz), theta (4 – 9 Hz), sigma (9 – 15 Hz), beta (15 – 30 Hz), low gamma (30 – 45 Hz) and high gamma (75 – 115 Hz) frequency bands.

Cortical coherence (a measure of undirected functional connectivity) between frontal and occipital channels was quantified by the multitaper time-frequency coherence method in Chronux analysis toolbox (Mitra and Bokil, 2008), with the same parameters described above for the spectral power analysis. The frontal-occipital coherence as a function of frequency was thus obtained by averaging the coherence over time in each behavioral state, for each mouse and treatment group. Coherence values were normalized by the Fisher’s z-transform (Miranda de Sá et al., 2009). The same frequency bands used in the spectral power analysis were assessed in the coherence analysis.

#### Analysis of directed connectivity

We used normalized symbolic transfer entropy (NSTE) to assess directed connectivity between frontal and occipital cortices. NSTE is an information theoretic measure, and our previous studies have validated its use to measure cortical connectivity changes in humans (Lee et al., 2013) and rats (Borjigin et al., 2013; Pal et al., 2016; Pal et al., 2020). In the calculation of NSTE, three parameters are required: embedding dimension (*d*_*E*_), time delay (τ), and transfer time (δ). We filtered the frontal and occipital EEG signals into the frequency bands as described above and segmented the filtered data into non-overlapped 5-s windows. We followed the methods used in previous studies (Li et al., 2017; Pal et al., 2020) and fixed the embedding dimension *d*_*E*_ = 3; as the time delay τ defines a broad frequency-specific window of sensitivity for NSTE (Jordan et al., 2013; Sitt et al., 2014; Ranft et al., 2016), we used τ = 64, 28, 17, 9, 6, 2, corresponding to delta (0.5 to 4 Hz), theta, sigma, beta, low- and high-gamma, respectively. For each window, we searched the transfer time δ = 1-50 (corresponding to 2 – 100 ms approximately) and selected the one that generated maximum feedback (from frontal to occipital) and feedforward (from occipital to frontal) NSTE, respectively. For statistical comparisons, the averaged connectivity values were calculated over the studied behavioral states for each mouse and treatment group.

#### Complexity analysis

We employed Lempel-Ziv Complexity (LZC) to quantify the dynamic changes in EEG signals from frontal and occipital cortices in the different behavioral states between the two treatment conditions. LZC is a method of symbolic sequence analysis (Lempel and Ziv, 1976) that has been shown to be a valuable tool to investigate the temporal or spatiotemporal complexity of brain activity (Schartner et al., 2017; Li and Mashour, 2019; Brito et al., 2020; Pal et al., 2020). Given the limited number of EEG channels in this study, we assessed only temporal complexity in frontal and occipital channels; spatiotemporal complexity was not evaluated. EEG signals were detrended using local linear regression (locdetrend function in Chronux analysis software) and a lowpass filter was applied at 115 Hz (but excluding the frequencies at 45-75Hz) via butterworth filter of order 4 (butter and filtfilt functions in MATLAB signal processing toolbox). The Hilbert transform of the signal was used to calculate the instantaneous amplitude, which was then segmented into 5-s windows without overlapping and binarized using its mean value as the threshold for each channel (Schartner et al., 2017). LZC analysis estimates the complexity of a finite series of numbers by computing the number of times that a different subsequence of consecutive characters, or “word,” is found within that series. To assess the signal complexity beyond the spectral changes, we generated surrogate data through phase randomization, while preserving the spectral profiles of the signal (Schartner et al., 2015; Schartner et al., 2017; Brito et al., 2020; Pal et al., 2020), and normalized the original LZC by the mean of the LZC values from N = 50 surrogate time series for each recording. The resultant LZC values were then averaged across all the windows as the estimate of the complexity for wakefulness, NREM sleep, and REM sleep. Higher values of LZC reflect higher complexity of the EEG signal. Analysis of spectral power, connectivity, and complexity during REM sleep was precluded by the profound suppression of REM sleep in the CNO group, which lead to an insufficient sample size.

### Immunohistochemistry and histological verification of designer receptor expression

After the last sleep experiment, mice were deeply anesthetized with isoflurane and perfused transcardially with ice-cold 0.1 M phosphate buffered saline pH 7.4 (PBS), followed by 5% formalin for 10 minutes using a MasterFlex perfusion pump (Cole Palmer, Vernon Hills, IL, USA). After perfusion, the brains were removed and post-fixed in 5% formalin overnight at 4°C. Subsequently, brains were cryoprotected with 20% sucrose in PBS for 1 to 2 days, and then transferred to 30% sucrose for an additional 2 to 3 days. Thereafter, brains were frozen in Tissue–Plus (Fisher Healthcare, Houston, TX, USA) and sectioned coronally at 40 μm using a cryostat (CM3050S, Leica Microsystems, Nussloch, Germany). Brain sections that contained the ventrolateral preoptic nucleus were collected and blocked in PBS containing 0.25% Triton X-100 and 3% normal donkey serum (Vector Laboratories, Burlingame, CA, USA) for 60 min at room temperature. Thereafter, sections were immunolabeled for mCherry. First, we incubated the tissues overnight at room temperature in primary antiserum (rat monoclonal anti-mCherry 1:30000; Thermo Fisher Scientific, Cat.#M11217). Next, sections were washed in PBS and incubated for 2 hours at room temperature in a donkey anti-rat secondary antiserum (1:500; Alexa Fluor 594, Thermo Fisher Scientific, Cat.#: A-21209). Last, brain sections were washed with PBS, float-mounted on glass slides and coverslipped with SlowFade Diamond (S36972 or S36973; Thermo Fisher Scientific). Brain sections containing the target region were then examined by means of fluorescence microscopy (BX43, Olympus America Inc., Waltham, MA, USA). The brain regions in which the designer receptor hM3Dq were expressed were identified with the aid of a mouse brain atlas (Paxinos and K.B.J., 2001). For stimulation experiments, only those mice that had reliable expression of designer receptors within the ventrolateral preoptic nucleus and adjacent areas were included in the analysis. Mice that received a vector injection but did not express designer receptors were used as controls.

### Statistical analyses

Statistical comparisons were performed with PRISM v7.0 (GraphPad Software Inc., La Jolla, CA). All data were tested for normality and are presented as mean ± standard error of the mean (SEM). A p < 0.05 was considered statistically significant. The effect of stimulation of preoptic glutamatergic neurons on total state duration, number of bouts, bout duration and number of NREM–to–W transitions over the 6-hour recording period was assessed by a one-tailed paired *t*-test. The directional hypothesis that predicted the nature of the effects of chemogenetic stimulation of preoptic glutamatergic neurons on these sleep-wake parameters was based on previously published data (Vanini et al., 2020). Differences in sleep-wake parameters for each 1-hour block were evaluated by a repeated-measures two-way analysis of variance (ANOVA) followed by a Bonferroni multiple comparisons test. The effect of CNO injection on the latency to NREM sleep and REM sleep onset was analyzed by a one-tailed paired *t*-test and by survival analysis. The effect of CNO administration on core body temperature was assessed by a one-way ANOVA. The treatment effect on EEG spectral power, Z’coherence, and NSTE (connectivity measures) for each frequency band was also assessed by a repeated-measures two-way ANOVA followed by a Bonferroni multiple comparisons test. Changes in LZC were analyzed by a two-tailed paired *t*-test. The effect of CNO administration on the FI over the 6-hour recording period was determined by a Wilcoxon signed-rank test; the Benjamini-Hochberg correction for multiple comparisons with a false discovery rate of 0.05 was used for the hour–by–hour analysis. Differences between treatment conditions in the distribution histograms were compared by the Kolmogorov-Smirnov test. Last, the relationship between REM sleep time (averaged for each mouse across the 6-hour recording period) and the fragmentation index or the number of NREM-wake transitions was estimated using the Spearman correlation coefficient.

## Results

### Anatomical localization of designer receptors

As described in our previous publication (Vanini et al., 2020), histological examination of vector injection sites revealed that neurons that expressed hM3Dq receptors were localized within the ventrolateral preoptic nucleus and the adjacent region encompassing (from medial to lateral) the ventral aspect of the medial preoptic nucleus as well as medial and lateral preoptic area (Figure 1). As stated above, in the current study we refer to this region collectively as medial-lateral preoptic. There was no correlation between the area of receptor expression and the stimulation effects on sleep-wake parameters. Importantly, there was no receptor expression in glutamatergic neurons, nor in fibers, within any structure of the basal forebrain region (i.e., horizontal limb of the diagonal band of Broca and substantia innominata), just lateral to the preoptic region of the hypothalamus.

**Figure 1.**
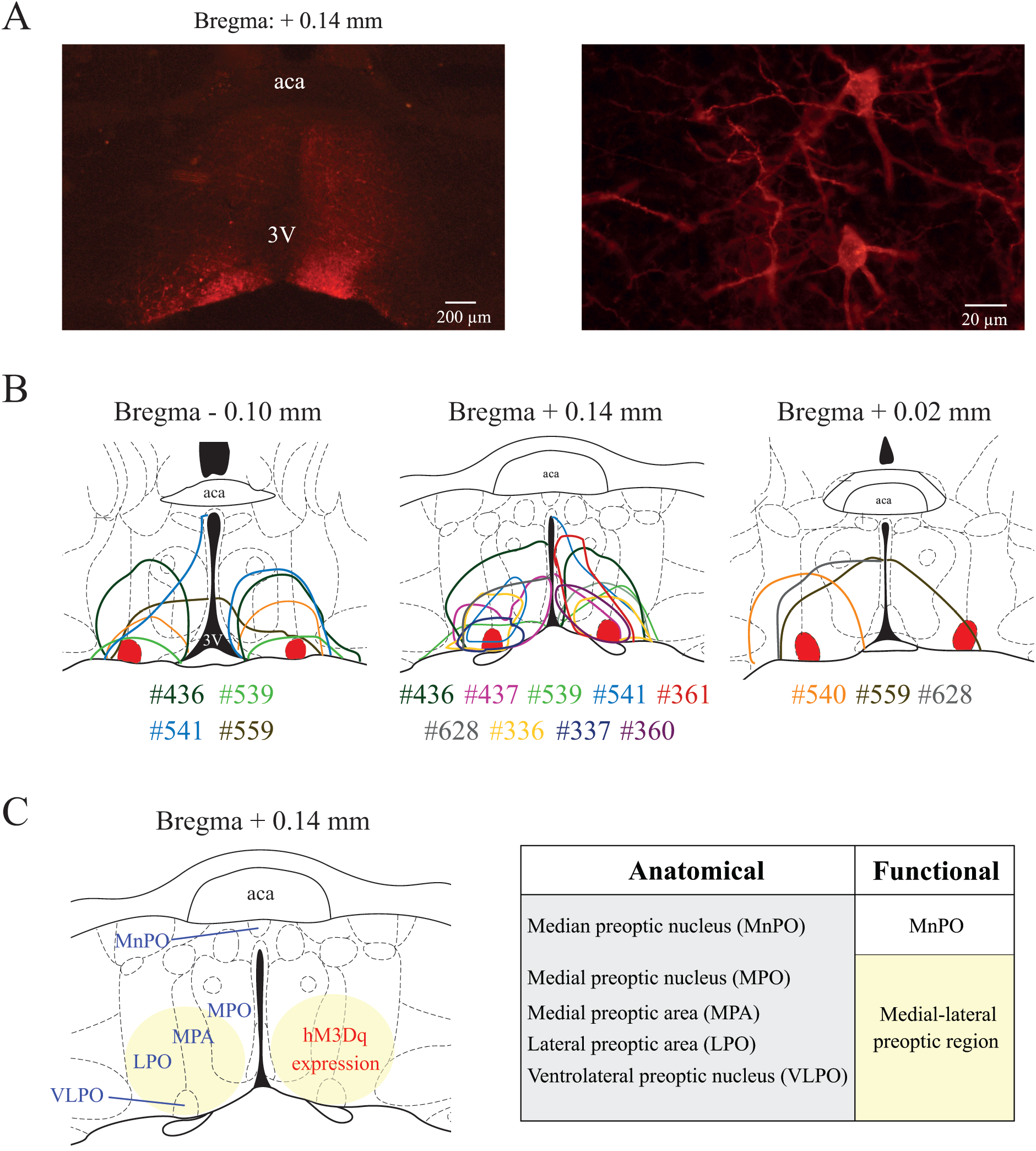
Histological confirmation of hM3Dq receptor expression in the preoptic area. *A.* The left and right panels show, respectively, examples of low and high magnification photographs of mCherry immunohistochemical staining (red) indicating the expression of the excitatory designer receptor hM3Dq within the ventrolateral preoptic nucleus of a Vglut2-Cre mouse. *B.* Color-coded depiction of the area of hM3Dq receptor expression is represented on coronal schematics of the preoptic area modified from a mouse brain atlas (Paxinos and K.B.J., 2001). For anatomical reference, the bilateral sites corresponding to the ventrolateral preoptic nucleus are marked in solid red. Color-matched identification numbers for each mouse used in this study are listed below each panel. *C.* Schematic of the preoptic area illustrating relevant anatomical subdivisions and main area of designer receptor expression (yellow circle). The chart below compares the anatomical subdivisions and functional nomenclature used in this study. Based on the uniform response to chemogenetic stimulation, different to that one observed in MnPO studies (Vanini et al., 2020), the area of hM3Dq receptor expression in the current study is referred as medial-lateral preoptic region. Abbreviations: aca, anterior commissure; 3V, third ventricle.

### Activation of medial-lateral preoptic glutamatergic neurons increased wakefulness and reduced NREM and REM sleep

Previously published data validated the viral vector for stimulation of preoptic neurons and showed that activation of preoptic glutamatergic neurons causes a robust increase in wakefulness (Vanini et al., 2020). The present study used a chemogenetic stimulation method and thorough analysis to better understand the role of these neurons in the regulation of behavioral states, sleep-wake state transitions, and cortical dynamics. Figure 2 illustrates the sleep-wake architecture in each Vglut2-Cre mouse (n = 11) expressing excitatory hM3Dq receptors in glutamatergic neurons of the medial-lateral preoptic region, after injection of vehicle or CNO. Figure 3 summarizes the effect of stimulation of preoptic glutamatergic neurons on sleep-wake parameters (time spent in wakefulness, NREM sleep and REM sleep, as well as the mean number and duration of bouts for each state) averaged across the 6-hour post-injection period. Compared to vehicle, CNO administration significantly reduced the time spent in REM sleep [t(10) = 6.72; p = 0.0094] (Figure 3A); this state was completely abolished in 6 out of the 11 mice studied (Figure 2). There was no significant difference in the time spent in wakefulness [t(10) = 0.58; p = 0.2870] and NREM sleep [t(10) =1.27; p = 0.1166]. Activation of preoptic glutamatergic neurons significantly increased the number of wakefulness and NREM sleep bouts [t(10) = 2.58; p = 0.0136 and t(10) = 2.53; p = 0.0149, respectively], and decreased the number of REM bouts [t(10) = 4.99; p = 0.0003] (Figure 3B). Furthermore, the duration of NREM sleep bouts was significantly reduced during the 6-hour recording period following CNO injection [t(10) = 2.56; p = 0.0141]; there were no changes in wakefulness bout duration [t(10) = 0.65; p = 0.2656] (Figure 3C). Because of the reduced number of mice that had REM sleep after CNO administration and the scarcity of REM sleep bouts in this group, the treatment effect on REM sleep bout duration was not statistically analyzed (vehicle vs. CNO: 13.98 ± 0.86 vs. 10.64 ± 2.75; Figure 3C).

**Figure 2.**
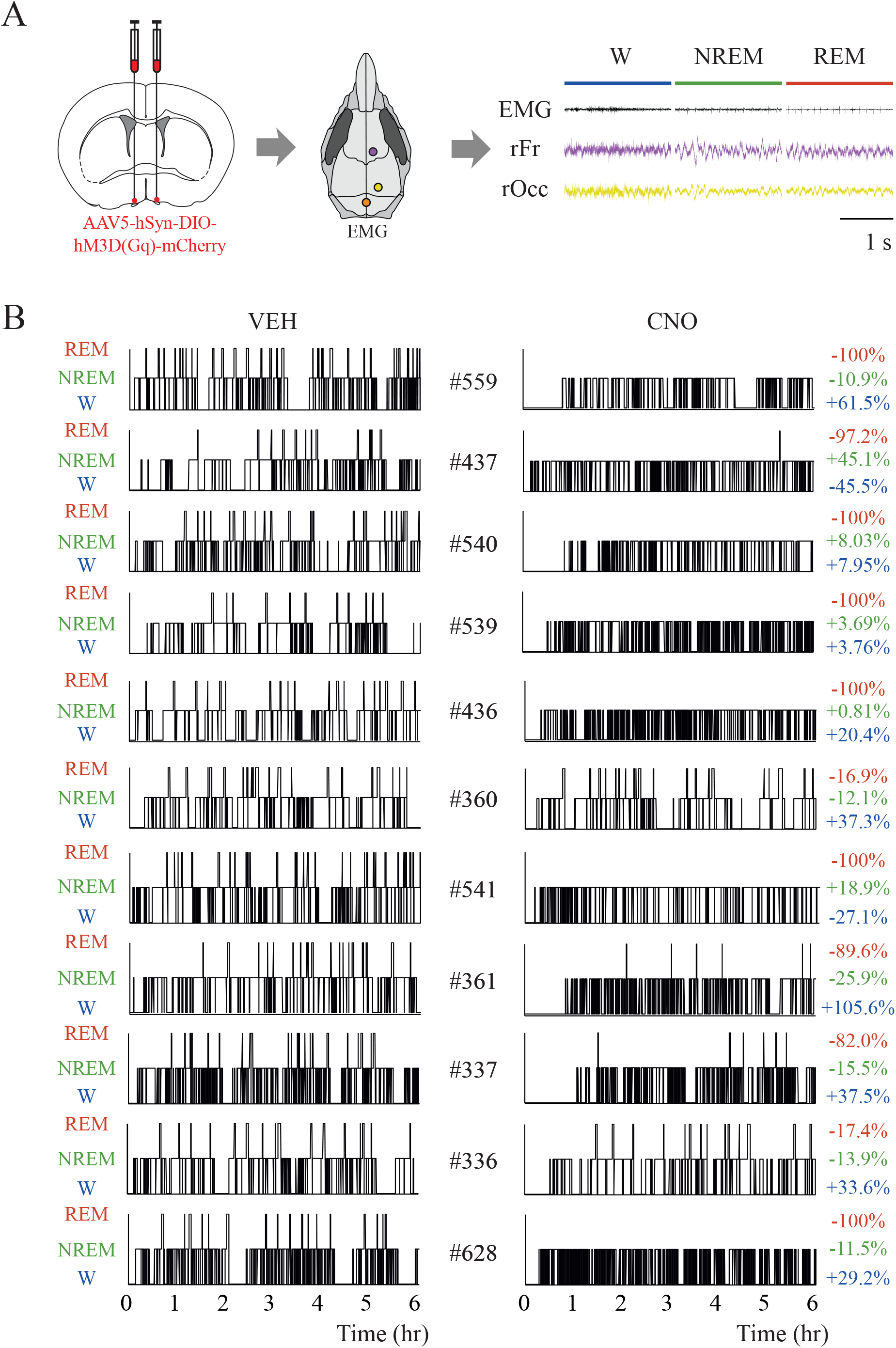
Chemogenetic activation of glutamatergic neurons in the medial-lateral preoptic region altered sleep-wake patterns. *A.* Schematic representation of bilateral injections of a Cre-dependent adeno-associated virus for expression of the excitatory designer receptor hM3Dq into the medial-lateral preoptic region of Vglut2-Cre mice. Three weeks after the injection, mice were implanted with electrodes for recording the electroencephalogram (EEG) from the right frontal (purple) and right occipital (yellow) cortex. A reference electrode was placed over the cerebellum (orange), and two electrodes were also implanted bilaterally in the neck muscles for recording the electromyogram (EMG). The right panel shows representative EEG and EMG signals from a mouse during wakefulness, NREM sleep and REM sleep. *B.* Hypnogram pairs illustrate the temporal organization of sleep-wake states after vehicle (VEH; left panels) and clozapine-N-oxide (CNO; right panels) for each mouse. The height of the bars (from lowest to highest) represents the occurrence of wakefulness (W), NREM sleep and REM sleep. Time 0 on the abscissa indicates the time at which the mouse received the injection of vehicle or CNO. Mouse identification numbers are listed between each pair of hypnograms. Individual differences relative to control (% change) in the time spent in wakefulness (blue), NREM sleep (green) and REM sleep (red) after CNO injection are shown to the right of each CNO hypnogram.

**Figure 3.**
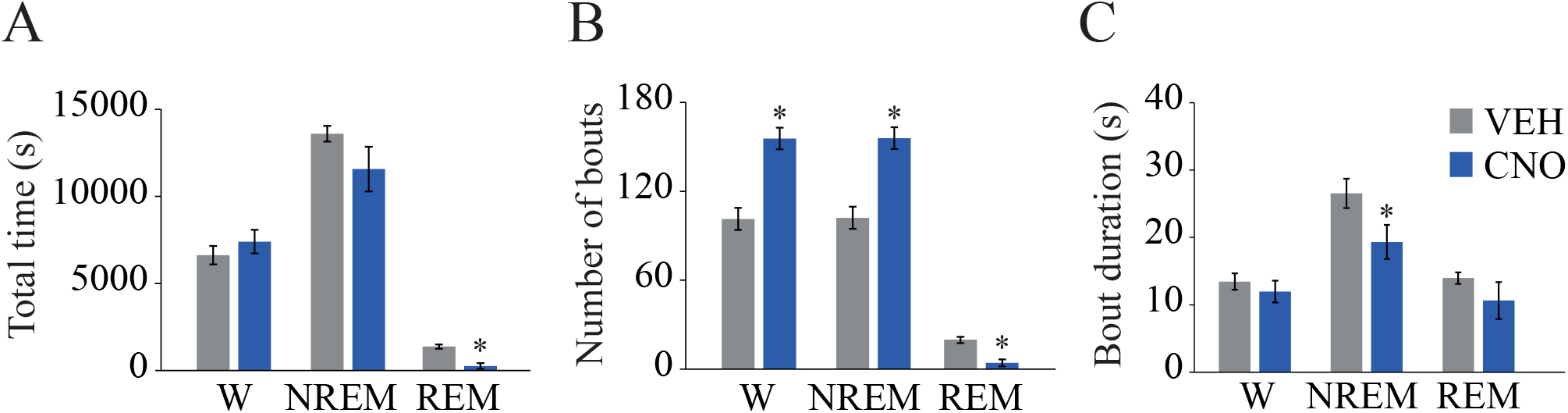
Mean effect of chemogenetic activation of glutamatergic neurons in the medial-lateral preoptic on sleep-wake states across the six-hour recording period. *A.* Changes in the time spent in wakefulness, NREM sleep and REM sleep. *B.* Effect on the number of wakefulness, NREM, and REM sleep bouts. *C.* Changes in the mean duration of wake, NREM, and REM sleep bouts. Data are reported as the mean ± standard error of the mean. One-tailed paired t-test was employed for statistical comparisons between treatment conditions. Asterisks indicate significant differences (p < 0.05) relative to control.

Figure 4A illustrates hour-by-hour changes in the time spent in wakefulness, NREM, and REM sleep as a function of treatment condition during the 6 hours post-injection. Repeated measures, two-way ANOVA indicated a significant treatment condition by time interaction for wakefulness [F(5,50) = 4.98; p = 0.0001] and NREM sleep [F(5,50) = 5.94; p = 0.0004]. CNO administration caused a significant increase in the time in wakefulness during the first hour post-injection (p = 0.0014) and a reduction of NREM sleep duration during hours 1 (p < 0.0001) and 2 (p = 0.0188) post-injection. For REM sleep, ANOVA revealed a significant drug effect [F(1,10) =45.12; p < 0.0001] and treatment by time interaction [F(5,50) = 2.69; p = 0.0316]. Activation of preoptic glutamatergic neurons caused a significant decrease in the time spent in REM sleep between hours 2 to 6 post-CNO injection (p = 0.002, < 0.0001, < 0.0001, < 0.0001 and < 0.0001). Figure 4B shows that, relative to control, CNO altered the number of wakefulness, NREM, and REM sleep bouts. ANOVA showed a significant drug effect on the number of wake (F(1,10) = 6.82; p = 0.0260) and NREM sleep bouts [F(1,10) = 6.29; p = 0.0311], and no significant effect on treatment condition by time interaction [F(5,50) = 1.41; p = 0.2380 and F(5,50) = 1.62; p = 0.1725]. The hour-by-hour post hoc analysis revealed that CNO significantly increased the number of wake (p = 0.0103, 0.0036, 0.0013 and 0.0048) and NREM sleep (p = 0.0129, 0.0033, 0.0013 and 0.0033) bouts during hours 2, 3, 4, and 6 post-injection. Additionally, there was a significant effect of CNO administration on the number of REM sleep bouts [F(1,10) = 26.43; p = 0.0004] and a significant treatment by time interaction [F(5,50) = 2.55; p = 0.0391]. Congruent with the decrease in REM sleep time, the mean number of REM sleep bouts was significantly reduced between hours 2 to 6 post-CNO administration (p <0.0001, <0.0001, <0.0001, <0.0001 and <0.0001). Figure 4C plots the mean bout duration for wakefulness, NREM sleep, and REM sleep. CNO significantly increased the duration of wake bouts (p = 0.0067) during the first hour post-injection, which accounts for the increase in the time in wakefulness shown in Figure 4A. Of note, the large deviation in the mean episode duration is related to one mouse that remained awake for the entire hour post-injection; removal of its vehicle and CNO data points confirmed that statistical significance was not driven exclusively by this mouse. NREM sleep episode duration was significantly decreased during hours 1 (p = 0.0010) and 2 (p = 0.0006) post-CNO injection. There were no significant changes in REM sleep episode duration.

**Figure 4.**
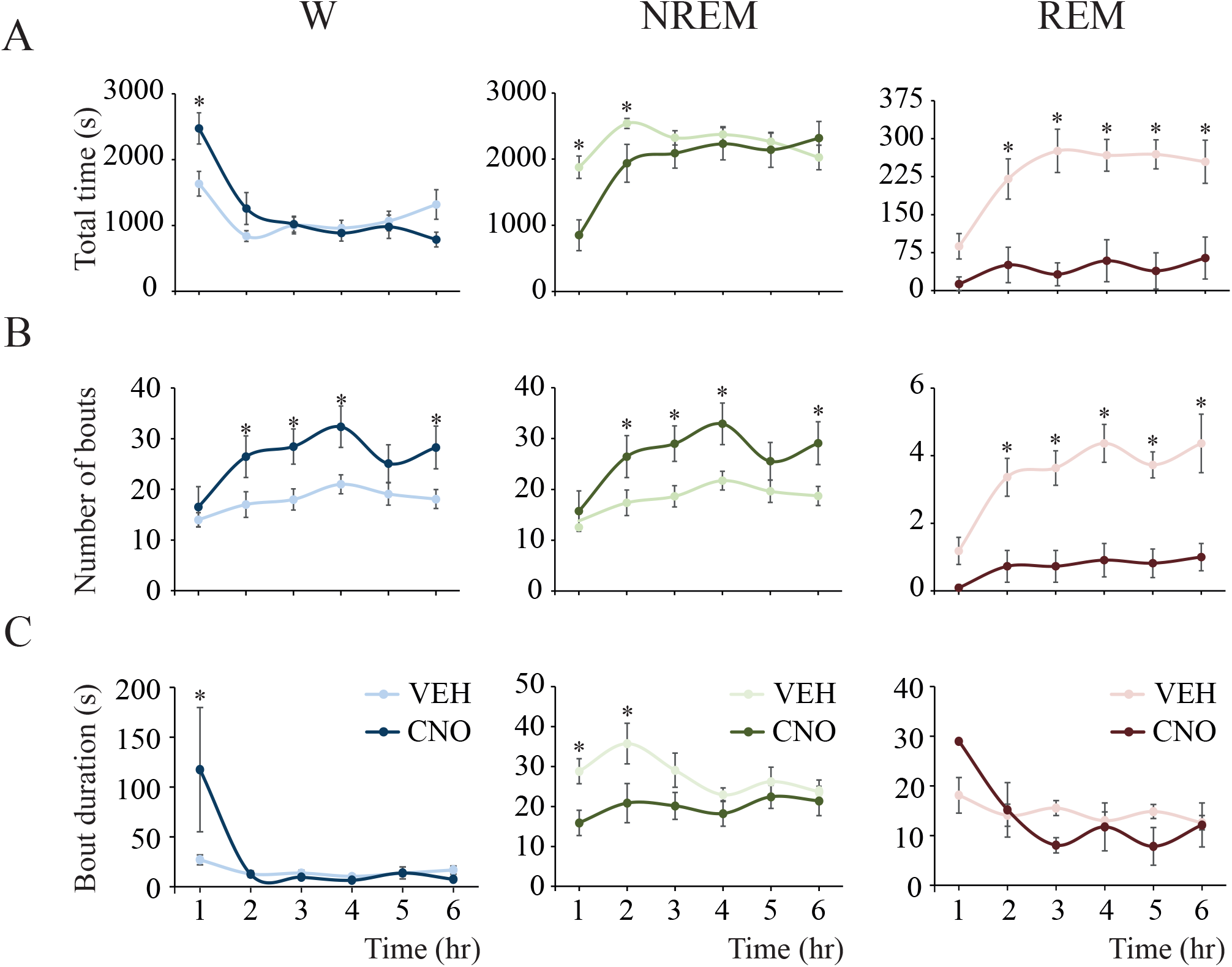
Activation of glutamatergic neurons in the medial-lateral preoptic region increased wakefulness and decreased NREM and REM sleep. Analyses of sleep-wake parameters were conducted in 1-hour blocks after vehicle (VEH; lighter color) and clozapine-N-oxide (CNO; darker color) injection. *A.* Time in wakefulness, NREM sleep and REM sleep. *B.* Number of bouts per state. *C.* Mean duration of wake, NREM and REM sleep bouts. Data are reported as the mean ± standard error of the mean. Two-way repeated measures ANOVA followed by a Sidak test adjusted for multiple comparisons was used to statistically compare sleep-wake parameters as a function of time and treatment condition. Asterisks indicate significant differences (p < 0.05) relative to control.

### Activation of medial-lateral preoptic glutamatergic neurons increased NREM and REM sleep latency

Figure 5 illustrates sleep latencies measured after vehicle and CNO injection. The graphs in 5A plot NREM and REM sleep latencies averaged across the 6-hour recording period for all mice. Animals that did not have REM sleep after CNO administration (n = 6) were assigned the maximum possible time (total recording time = 21600 s) as the latency to REM sleep. Relative to vehicle, CNO injection significantly increased the latency to both NREM [Mean ± SEM = 604.55 ± 102.51 vs 1796.36 ± 378.39, t(10) = 2.88; p = 0.0081] and REM sleep [Mean ± SEM = 3334.55 ± 494.74 vs 15419.55 ± 2463.17, t(10) = 4.76; p = 0.0004]. Because several mice did not have REM sleep after CNO injection, changes in the latency to the first REM sleep bout (also the latency to NREM sleep onset) was assessed by survival analysis (Vanini and Baghdoyan, 2013). The graphs in 5B plot the probability of not having NREM sleep and REM sleep after CNO administration. Activation of preoptic glutamatergic neurons significantly increased the probability of not having NREM sleep (p = 0.0006) and REM sleep (p = 0.0001). Furthermore, relative to control, the probability of REM sleep not occurring after CNO administration remained elevated (> 50%) until the end of the recording period.

**Figure 5.**
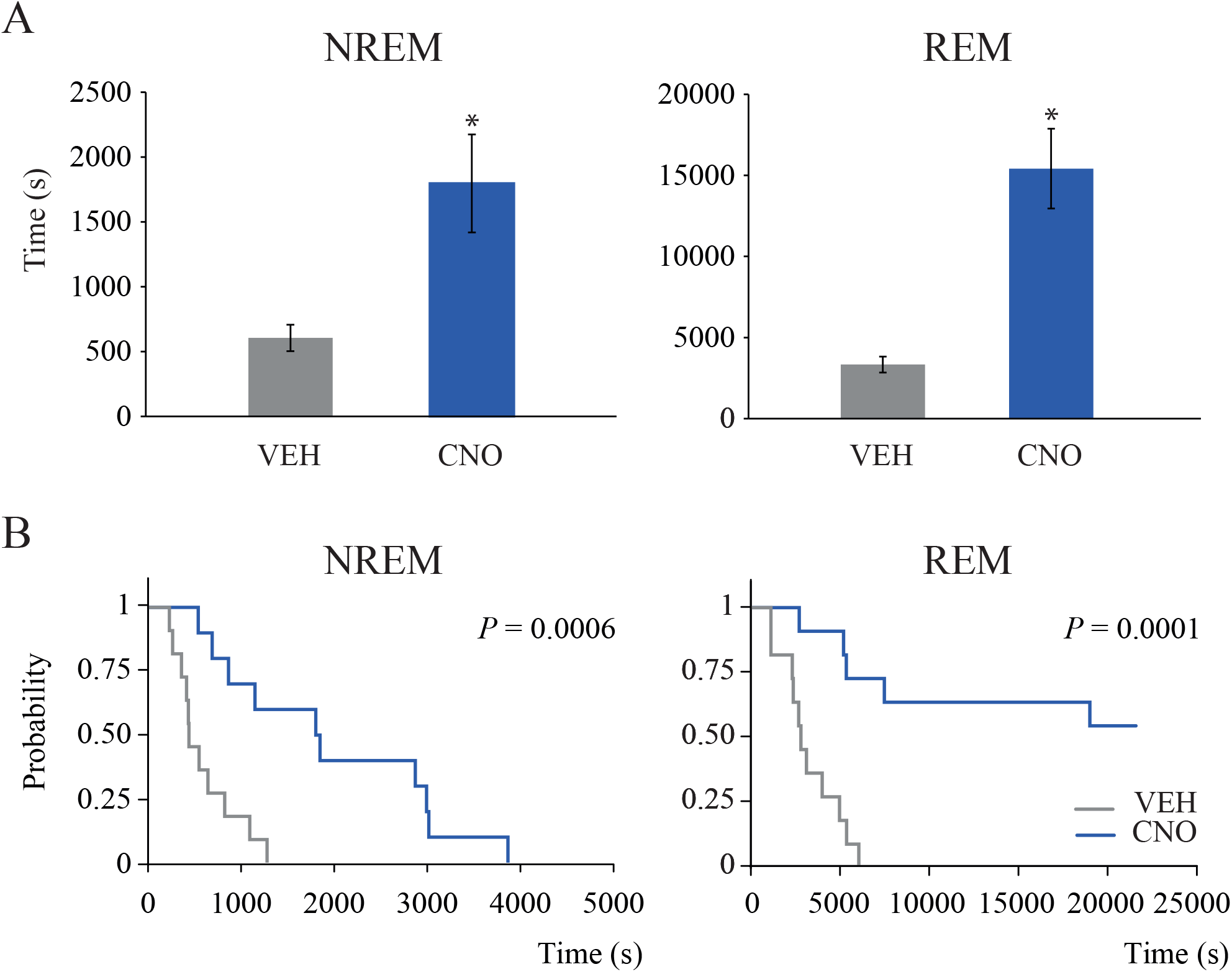
Activation of glutamatergic neurons in the medial-lateral preoptic region increased the latency to NREM and REM sleep. *A.* Effect of clozapine-N-oxide (CNO) injection on NREM (left) and REM sleep (right) latencies. Data are reported as the mean ± standard error of the mean. One-tailed paired t-test was employed for statistical comparisons. Asterisks indicate significant differences (p < 0.05) relative to control. *B.* The graphs show the probability of no NREM sleep (left) and REM sleep (right) generation after injection of vehicle (VEH) or CNO. Survival analysis demonstrated that the probability of no REM sleep occurring remained increased throughout the 6-hour recording period and was significantly different between treatment condition in both NREM (p = 0.0006) and REM sleep (p = 0.0001).

### Activation of medial-lateral preoptic glutamatergic neurons increased NREM to Wake transitions

Because activation of preoptic glutamatergic neurons increased the number of wake and NREM sleep bouts, we compared the number of transitions from NREM sleep to wakefulness as a function of treatment condition and time. Figure 6A plots the mean number of NREM to wake transitions averaged across the 6-hour recording period for all mice. CNO injection significantly increased the number of transitions by 76% [t(10) = 3.36; p = 0.0036]. Figure 6B illustrates hour–by–hour changes in NREM to wake transitions during 6 hours after vehicle and drug administration. ANOVA revealed a significant effect of treatment (F(1,10) = 11.35; p = 0.0071). There was no treatment by time interaction [F(5,50) = 1.96; p = 0.1012]. Multiple comparisons post hoc tests showed that the number of transitions was significantly increased between post-injection hours 2 to 6 (p = 0.0014, 0.0001, <0.0001, 0.0168 and 0.0002). There was no significant correlation between REM sleep time and the number of NREM to wake transitions (r =−0.4808; p = 0.1364).

**Figure 6.**
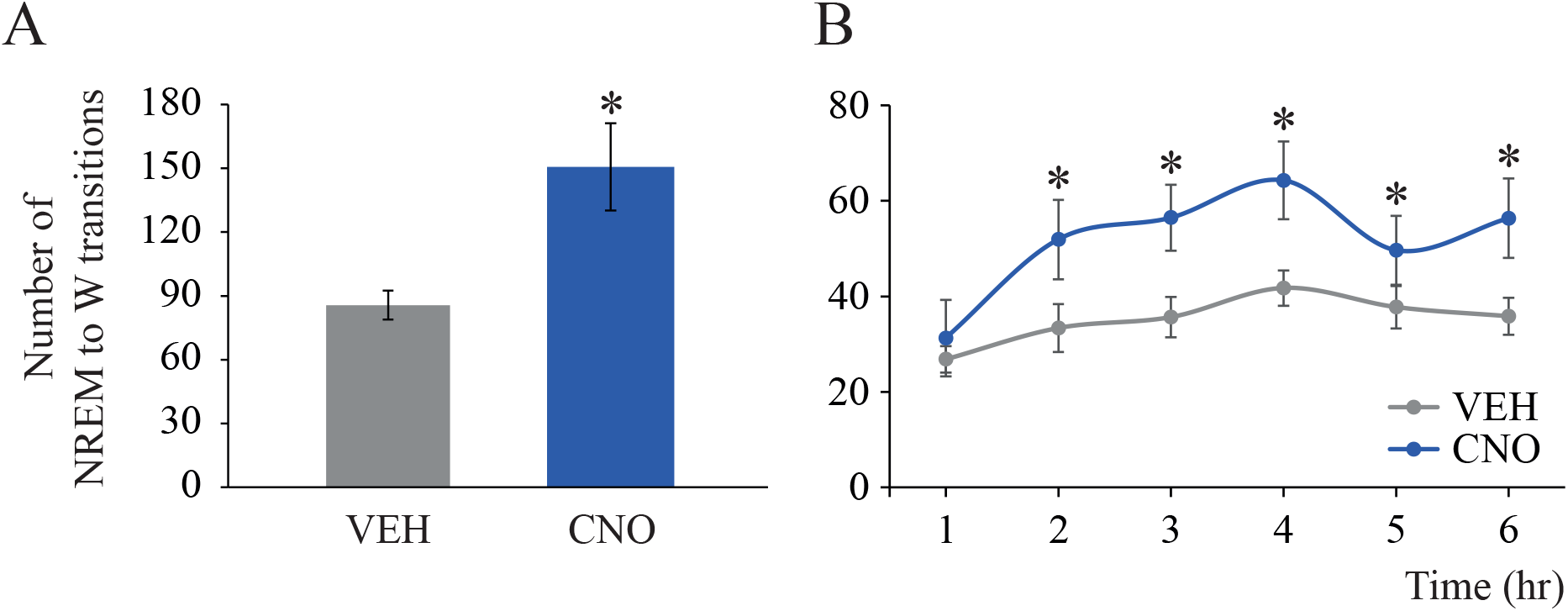
Activation of glutamatergic neurons in the medial-lateral preoptic region increased NREM to wake transitions. Number of transitions from NREM to wakefulness (W) averaged across the 6-hour recording period (*A*) and per 1-hour block (*B*) after injection of vehicle (VEH) or clozapine-N-oxide (CNO). Data are reported as the mean ± standard error of the mean. One-tailed paired t-test (6-hour block) and two-way repeated measures ANOVA followed by a Sidak test corrected for multiple comparisons (1-hour block analysis) was used for statistical comparisons between treatment conditions. Asterisks indicate significant differences (p < 0.05) relative to control.

### Activation of medial-lateral preoptic glutamatergic neurons caused a robust NREM sleep fragmentation

Based on (1) the increase in the number of NREM sleep bouts, (2) the reduction of NREM sleep bout duration, and (3) the increment in the number of NREM to wake transitions after CNO injection, we hypothesized that the activation of preoptic glutamatergic neurons causes NREM sleep instability. We tested this hypothesis using three different approaches.

First, we calculated the state transition probability after vehicle and CNO administration by means of a Markov model. Figure 7A depicts the probability of transitioning between/within states of wakefulness, NREM sleep and REM sleep. Consistent with a previous study (Perez-Atencio et al., 2018), the probability of remaining in one state was much higher than the probability of transitioning into a different state. Relative to vehicle, activation of glutamatergic neurons in the medial-lateral preoptic region significantly reduced the probability of remaining in NREM sleep [Mean ± SEM = 0.9384 ± 0.009 vs 0.9621 ± 0.003, *W* [*probability vector value*] =−7; p = 0.0186], while there was no significant change in the probability of remaining in wakefulness [Mean ± SEM = 0.8983 ± 0.0491 vs 0.9205 ± 0.0069, *W* =−24; p = 0.1602] or REM sleep [0.9052 ± 0.040 vs 0.9621 ± 0.003, *W* =−7; p = 0.2188]. Consistent with this evidence, there was an increased probability of transitioning from NREM sleep to wakefulness [Mean ± SEM = 0.060 ± 0.010 vs 0.031 ± 0.003, *W* = 64; p = 0.0010], while the probability of transitioning from wakefulness to NREM sleep was not affected [Mean ± SEM = 0.1017 ± 0.015 vs 0.079 ± 0.007, *W* = 24.0; p = 0.1602]. Furthermore, as expected because of the drastic reduction in the amount of REM sleep, CNO significantly decreased the probability of transitioning from NREM to REM sleep [Mean ± SEM = 0.0017 ± 0.001 vs 0.0072 ± 0.001, *W* =−60; p = 0.0024]. The probability of entering either wakefulness or NREM sleep from REM sleep was not affected by the activation of the glutamatergic neurons of the medial-lateral preoptic region [Mean ± SEM = 0.0833 ± 0.032 vs 0.0625 ± 0.008, *W* = 7; p = 0.2188 and 0.011 ± 0.009 vs 0.003 ± 0.001, *W* = 2; p = 0.3750, respectively]. None of the mice entered REM sleep from wakefulness after vehicle−as expected-or CNO administration.

**Figure 7.**
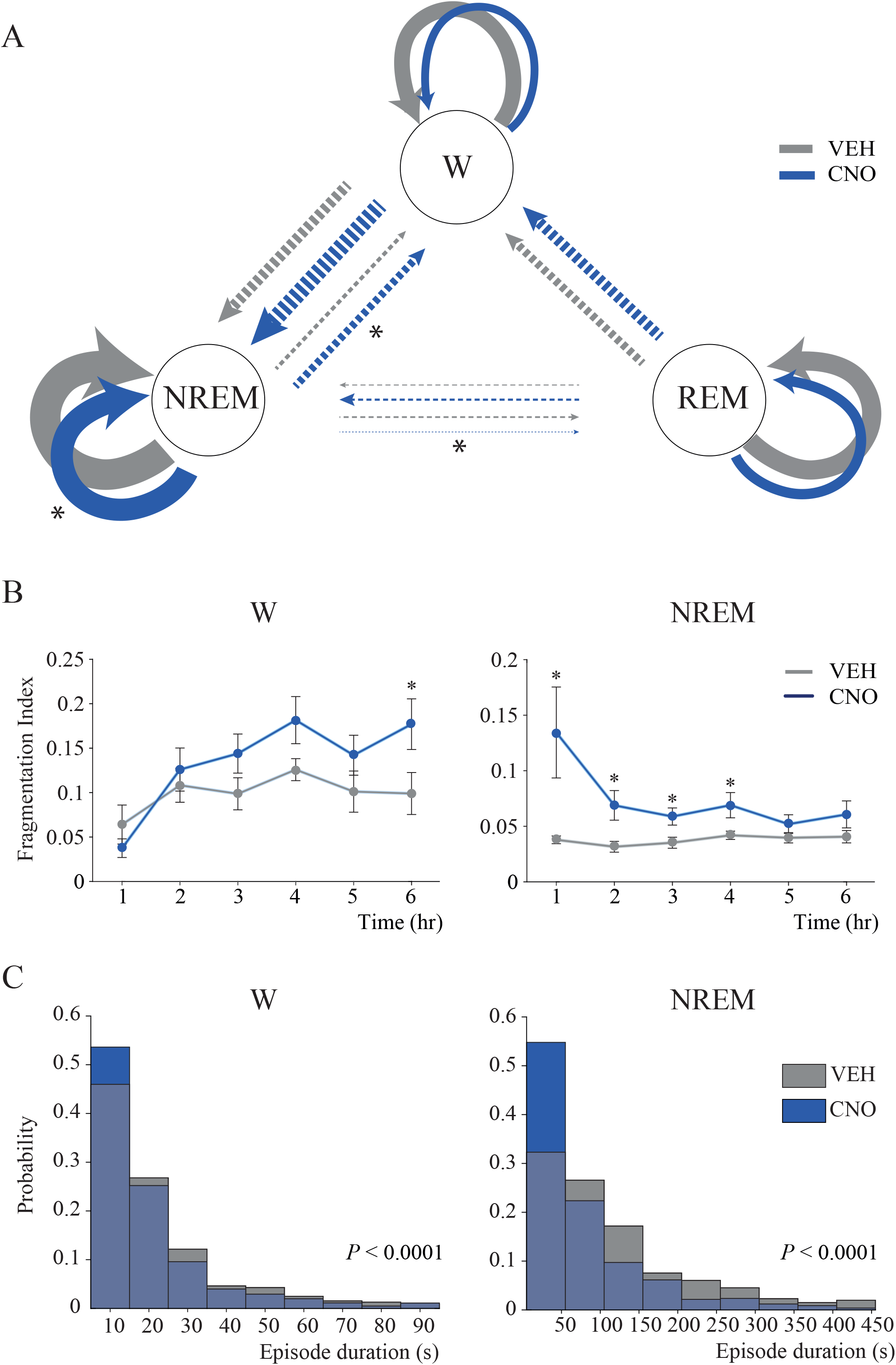
Activation of glutamatergic neurons in the medial-lateral preoptic region increased NREM sleep fragmentation. *A.* Diagram of the Markov Model for wakefulness (W) – NREM sleep – REM sleep dynamics after vehicle (VEH) and clozapine-N-oxide (CNO) injection. Circular arrows indicate the probability of remaining within the same state and straight arrows indicate the probability of transitioning from one state to another. The thickness of the arrows is proportional to the corresponding probability. Two different scales were used, one for the circular arrows and another one for straight arrows, i.e., circular arrows were designed with continuous lines while straight arrows were designed with dotted lines. Differences between vehicle and CNO were analyzed by means of Wilcoxon matched-pairs rank tests. Asterisks indicate significant differences (p < 0.05) relative to control. *B.* Fragmentation index (FI) for each 1-hour block during wakefulness and NREM sleep after vehicle or CNO injection. We defined FI as FI = 1 - p(X/X), being p(X/X) the probability of transitioning from the state X to the same state X. This implies that FI = 1 only if the state is completely fragmented, i.e., if the probability of transitioning to the same state equals to 0. Error bars represent the standard error of the mean. Differences between vehicle and CNO were analyzed by means of Wilcoxon matched-pairs rank tests and *p* values were corrected by the Benjamini and Hochberg for a false discovery rate of 5%. Asterisks indicate significant differences (p < 0.05) relative to control. *C.* Histograms illustrate episode durations in units of 10 s of wakefulness and in units of 50 s of NREM sleep. Vehicle and CNO histograms were overlapped to better appreciate the differences. The difference in the distribution of episode duration between vehicle and CNO injection was analyzed by means of a Kolmogorov-Smirnov test. Episode duration after CNO and vehicle had a different distribution in both wakefulness (p < 0.0001) and NREM sleep (p < 0.0001), with increased short and reduced long bouts after CNO administration.

Second, to evaluate the stability of sleep-wake states in each 1-hour block, we calculated a fragmentation index (FI); FI = 1 indicates that the state is maximally unstable and fragmented. Because of the scarcity of REM sleep after CNO administration, we only performed this analysis for wakefulness and NREM sleep states. Figure 7B shows that, in comparison to vehicle, stimulation of glutamatergic neurons in the medial-lateral preoptic region significantly fragmented NREM sleep during the first 4 hours of the recording period [*W* = 45, p = 0.0294; *W* = 54, p = 0.0294; *W* = 48, p = 0.0315; *W* = 46, p = 0.0315]. There was no significant correlation between REM sleep time and FI NREM values (r =−0.3569; p = 0.2785). Additionally, wakefulness was only fragmented during hour 6 post-injection [*W* = 64, p = 0.0060].

Third, to understand better the effect of the activation of preoptic glutamatergic neurons on sleep and wake episode duration, we compared the distribution of NREM sleep and wake bout duration after vehicle and CNO injection. Figure 7C shows that the activation of glutamatergic neurons of the medial-lateral preoptic region significantly altered the distribution of bout duration by increasing the number of short bouts and decreasing the number of long bouts during both NREM sleep (p < 0.0001) and wakefulness (p < 0.0001).

### Activation of medial-lateral preoptic glutamatergic neurons did not modify body temperature

Activation of MnPO glutamatergic (Abbott and Saper, 2017; Vanini et al., 2020) and VLPO galaninergic neurons (Kroeger et al., 2018) causes hypothermia in mice, which can increase wakefulness and reduce NREM and REM sleep duration (Parmeggiani, 1987). Thus, the effect of glutamatergic neurons in the medial-lateral preoptic region on core body temperature was evaluated pre- and post-CNO injection (Figure 8). The mean temperature decrease post-CNO injection (from min 10 to 90) was 0.50 ± 0.07 _ (mean ± SEM). A repeated measures one-way ANOVA confirmed that activation of these neurons did not significantly alter core body temperature [F(1.945, 7781) = 1.757; p = 0.2345].

**Figure 8.**
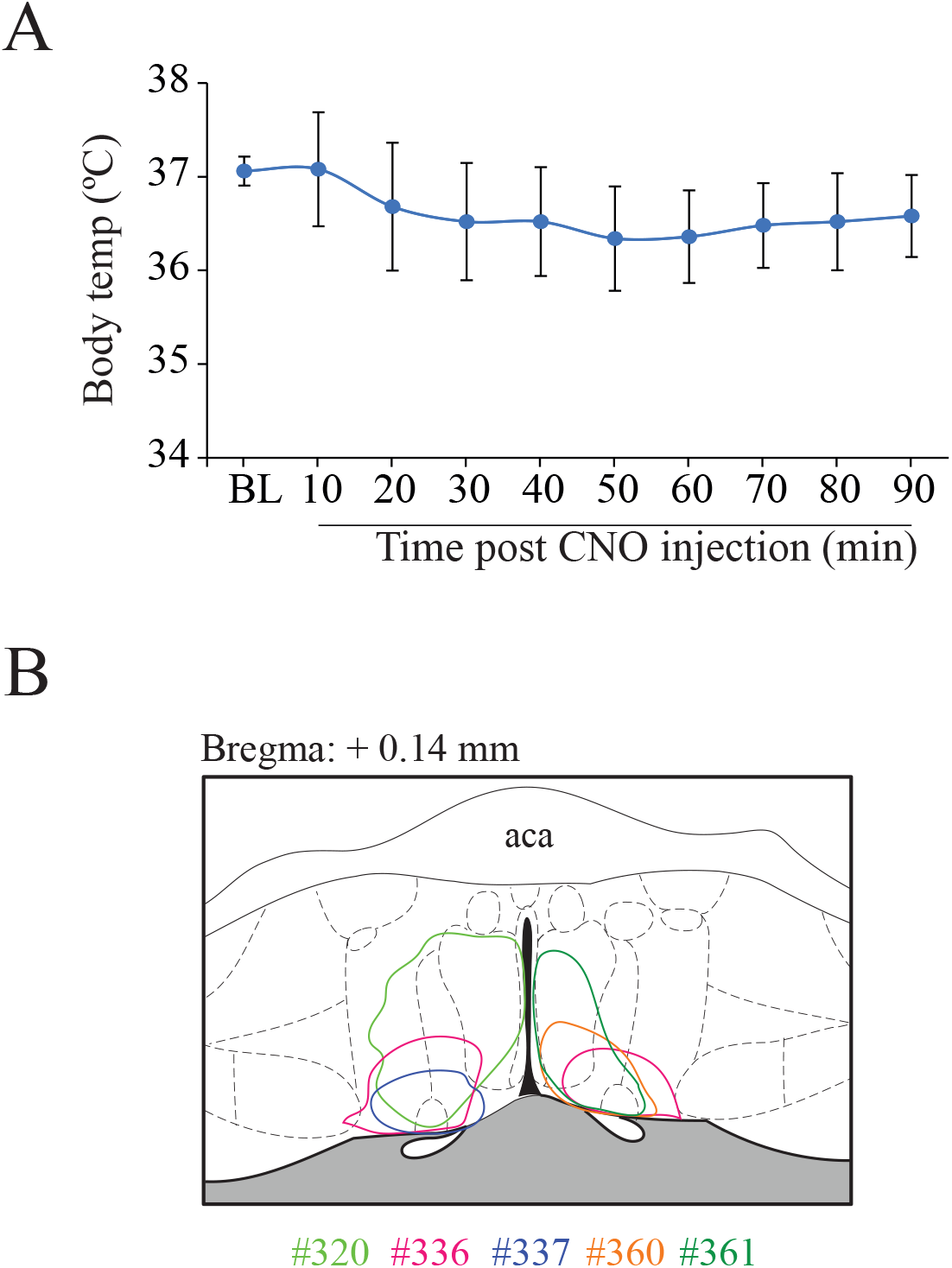
Activation of glutamatergic neurons in the medial-lateral preoptic region did not alter core body temperature. *A.* The graph illustrates the time course of core body temperature in awake mice after injection of clozapine-N-oxide (CNO) for activation of glutamatergic neurons within the medial-lateral preoptic region. Data are reported as the mean ± standard error of the mean. Repeated measures one-way ANOVA was used for statistical comparisons of mean temperature levels after CNO injection relative to baseline (BL). *B.* Color-coded area of hM3Dq receptor expression within the medial-lateral preoptic region of Vglut2-Cre mice used for temperature experiments, represented on a coronal schematic of the preoptic area modified from a mouse brain atlas (Paxinos and K.B.J., 2001).

### CNO administration to control mice without designer receptors did not alter sleep-wake states

To rule out any non-specific effect of CNO or its active metabolites (Ilg et al., 2018), as well as from the cell damage caused by the injection procedure or vector-associated toxicity (Rezai Amin et al., 2019) we included two negative control groups. The effect of 1.0 mg/kg CNO on sleep-wake states was evaluated in a group of mice that received the vector injection into the medial-lateral preoptic region but did not express hM3Dq receptors (n=5), and in a second group injected with the “empty” control vector that only contained the fluorescent reporter mCherry (n=4). Consistent with previous work (Vanini et al., 2020), CNO injection did not alter sleep-wake states in either group. Therefore, the data from all 9 mice were pooled for statistical analysis. The data corresponding to each control group (6-hour average) are shown in Table 1. Figure 9A-C summarizes group data averaged across the 6-hour recording period and shows that CNO injection did not modify the time in wakefulness [t(8) = 0.069; p = 0.9460], NREM sleep [t(8) = 0.351; p = 0.7344] and REM sleep [t(8) = 0.506; p = 0.6265]. Neither the number of bouts nor their duration was altered during wakefulness [t(8) = 1.182; p = 0.2710 and t(8) = 1.850; p = 0.1014], NREM sleep [t(8) = 1.209; p = 0.2613 and t(8) = 1.215; p = 0.2589] and REM sleep [t(8) = 0.525; p = 0.6137 and t(8) = 0.142; p = 0.8909]. Furthermore, there were no significant changes in the total time in wakefulness, NREM sleep, and REM sleep in the hour-by-hour analysis (Figure 9D-F). Repeated measures, two-way ANOVA showed no significant effect of the treatment, or treatment by time interaction in wakefulness [F(1,8) = 0.004; p = 0.9460 and F(5,40) = 0.124; p = 0.9860), NREM sleep (F(1,8) = 0.123; p = 0.7344 and F(5,40) = 0.054; p = 0.9980] and REM sleep [F(1,8) = 0.256; p = 0.6265 and F(5,40) = 2.173; p = 0.0763]. The latency to NREM or REM sleep was not significantly altered by CNO, as shown in Figure 9G [Mean ± SEM = 336.1 ± 143.0 vs 598.9 ± 142.0, t(8) = 1.986; p = 0.0823 and 3530 ± 1034 vs 4929 ± 2202, (t(8) = 0.662; p = 0.5268] respectively. Importantly, neither wakefulness nor NREM sleep were fragmented during the 6-hour period after CNO injection [Mean ± SEM = 0.075 ± 0.005 vs 0.068 ± 0.003, *W* =−27; p = 0.1289] and [Mean ± SEM = 0.046 ± 0.005 vs 0.040 ± 0.003, *W* =−21; p = 0.2500]. Accordingly, no fragmentation was found in any of the 6 one-hour block analyses for wakefulness [*W* =−27, p = 0.1289; *W* =−3.0, p = 0.9102; *W* =−11, p = 0.5703; *W* = 1.0, p = > 0.9999; *W* =−19, p = 0.3008; *W* = 1.0, p = > 0.9999) or NREM sleep (*W* =−15, p = 0.4258; *W* =−27, p = 0.1289; *W* =−11, p = 0.5703; *W* = 9.0, p = 0.6523; *W* =−13, p = 0.4961; *W* = 7.0, p = 0.7344] as shown in Figure 9I.

**Table 1.** Sleep-wake parameters obtained from individual, negative control groups as a function of treatment condition.

|  | AAV-mCherry<br>("empty" control vector) |  | AAV without hM3Dq<br>expression |  |
| --- | --- | --- | --- | --- |
|  | VEH | CNO | VEH | CNO |
| Time spent in Wake (s) | 7919 ± 633.5 | 7730 ± 353.3 | 7250 ± 484.3 | 7347 ± 435.4 |
| Time spent in NREM sleep (s) | 12443 ± 516.9 | 12321 ± 347.5 | 12803 ± 482.7 | 13166 ± 374.3 |
| Time spent in REM sleep (s) | 1239 ± 312.3 | 1549 ± 77.55 | 1547 ± 55.6 | 1087 ± 262.3 |
| Number of Wake bouts | 126.8 ± 17.54 | 109.3 ± 5.22 | 107.2 ± 15.68 | 100.0 ± 13.88 |
| Number of NREM sleep bouts | 126.8 ± 17.72 | 108.8 ± 5.28 | 107.8 ± 15.62 | 100.6 ± 14.09 |
| Number of REM sleep bouts | 17.50 ± 2.46 | 19.75 ± 3.42 | 20.6 ± 2.22 | 16.40 ± 3.95 |
| Duration of Wake bout (s) | 12.84 ± 1.01 | 14.30 ± 1.22 | 14.29 ± 1.63 | 15.36 ± 1.23 |
| Duration of NREM sleep bout (s) | 20.93 ± 3.13 | 22.75 ± 1.04 | 25.60 ± 3.40 | 28.56 ± 4.32 |
| Duration of REM sleep bout (s) | 14.18 ± 3.17 | 17.13 ± 2.84 | 15.52 ± 1.22 | 12.60 ± 1.70 |
| Latency to NREM sleep (s) | 356.3 ± 237.1 | 278.8 ± 139.6 | 320 ± 199.2 | 855 ± 154.7 |
| Latency to REM sleep (s) | 4256 ± 1993 | 1813 ± 125.6 | 2949 ± 1128 | 7423 ± 3735 |

**Figure 9.**
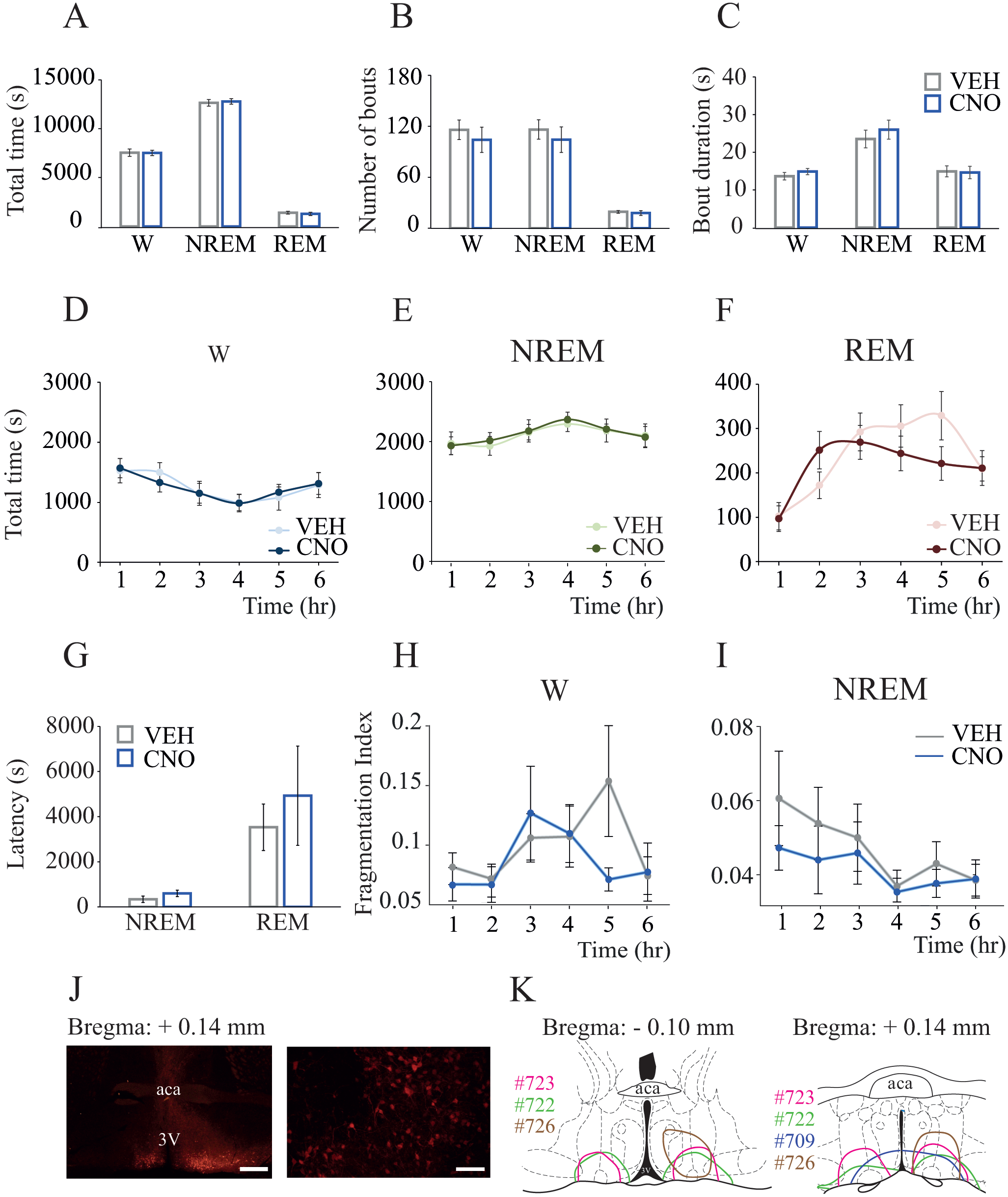
CNO administration to mice that did not express designer receptors did not alter sleep-wake states. Sleep data from mice injected with (1) AAV-hSyn-DIO-hM3D(Gq)-mCherry into the preoptic area but did not express designer receptors and (2) the control vector AAV-hSyn-DIO-mCherry were pooled together and used as negative controls. *A.* Group data summarizing total time in wakefulness, NREM sleep and REM sleep after vehicle (VEH) or clozapine-N-oxide (CNO) injection. *B.* Number of bouts of wakefulness, NREM and REM sleep. *C.* Comparison of mean episode duration per state. *D-F.* The graphs plot the mean time in wakefulness, NREM sleep and REM sleep, respectively, for each 1-hour block after vehicle and CNO administration. *G.* Latency to NREM and REM sleep. Data are reported as the mean ± standard error of the mean. A one-tailed paired t-test was used for statistical comparisons between treatment conditons. Asterisks indicate significant differences (p < 0.05) relative to control. *H and I.* Fragmentation index (FI) calculated for NREM and REM sleep, respectively, plotted for each 1-hour block after vehicle and CNO injection. Wilcoxon matched-pairs rank tests were used for statistical comparisons between treatment conditions. *J.* The left and right panels show, respectively, low and high magnification photographs of mCherry immunohistochemical staining (red) corresponding to the fluorescent reporter of the control vector within the ventrolateral preoptic nucleus of a Vglut2-Cre mouse. Calibration bars, 500 and 100 μm, respectively. *K.* Color-coded depiction of control vector injection area, represented on coronal schematics of the preoptic area modified from a mouse brain atlas (Paxinos and K.B.J., 2001). Color-matched identification numbers for each mouse used in this study are listed on the left side of each panel. Abbreviation: aca, anterior commissure; 3V, third ventricle.

### Activation of glutamatergic neurons in the MnPO did not substantially alter sleep-wake states

To test whether the effects of the activation of glutamatergic neurons in the medial-lateral preoptic region were site-specific, we reanalyzed sleep-wake data from a previous study in which we used the same chemogenetic methods to stimulate MnPO glutamatergic neurons in Vglut2-Cre mice (Vanini et al., 2020). Figure 10A-C shows that activation of glutamatergic neurons within the MnPO (n = 7 mice) did not modify the time in wakefulness [t(6) = 0.253; p = 0.8087], NREM sleep [t(6) = 0.748; p = 0.2412] and REM sleep [t(6) = 1.738; p = 0.066]. CNO administration did not alter the number of bouts or bout duration of wakefulness [t(6) = 0.577; p = 0.2925 and t(6) = 0.630; p = 0.5519], NREM [t(6) = 0.635; p = 0.2745 and t(6) = 0.223; p = 0.8312], and REM sleep [t(6) = 2.294; p = 0.0616 and t(6) = 0.889; p = 0.4082]. Additionally, there were no significant changes in the time spent in wake, NREM, and REM sleep in the 1-hour block analysis (Figure 10D-F). Repeated measures, two-way ANOVA revealed no significant effect of treatment or treatment by time interaction for wakefulness [F(1,6) = 0.06; p = 0.808 and F(5,30) = 0.37; p = 0.866], NREM sleep [F(1,6) = 0.56; p = 0.482 and F(5,30) = 0.24; p = 0.939] and REM sleep [F(1,6) = 3.23; p = 0.122 and F(5,30) = 0.85; p = 0.524]. Figure 10G shows that activation of glutamatergic neurons in the MnPO did not alter the latency to NREM [Mean ± SEM = 568.6 ± 136.5 vs 680.0 ± 199.0, t(6) = 0.619; p = 0.558] or REM sleep [2676 ± 330.3 vs 4496 ± 777.0, t(6) = 2.14; p = 0.076]. Furthermore, activation of MnPO glutamatergic neurons did not cause fragmentation of wake or NREM sleep states (Figure 10H-I).

**Figure 10.**
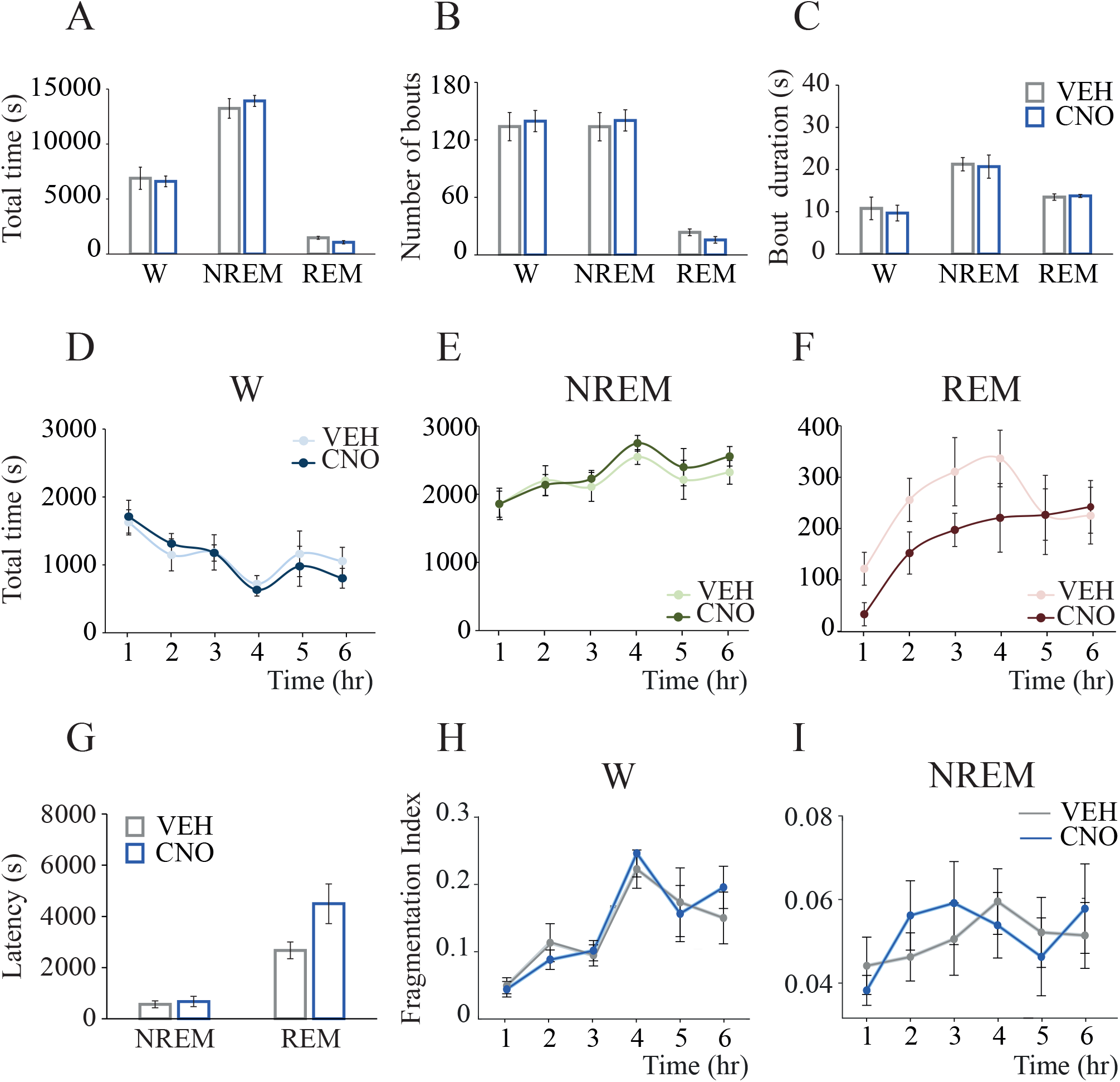
Activation of glutamatergic neurons in the Median Preoptic Nucleus (MnPO) of the hypothalamus did not substantially alter sleep-wake states. Sleep data from a previous study (collected using identical procedures and experiment design (Vanini et al., 2020)) were reanalyzed and used in the current study as a site-control group. All Vglut2-Cre mice included in this control group had confirmed hM3Dq receptor expression in MnPO glutamatergic neurons. *A.* Group data summarizing total time in wakefulness, NREM sleep and REM sleep after vehicle (VEH) or clozapine-N-oxide (CNO) injection. *B.* Number of bouts of wakefulness, NREM and REM sleep. *C.* Comparison of mean episode duration per state. *D-F.* The graphs plot the mean time in wakefulness, NREM sleep and REM sleep, respectively, for each 1-hour block after vehicle and CNO administration. *G.* Latency to NREM and REM sleep. Data are reported as the mean ± standard error of the mean. A one-tailed paired t-test was used for statistical comparisons. Asterisks indicate significant differences (p < 0.05) relative to control. *H and I.* Fragmentation index (FI) calculated for NREM and REM sleep, respectively, plotted for each 1-hour block after vehicle and CNO injection. Wilcoxon matched-pairs rank tests were used for statistical comparisons between treatment conditions.

### Activation of medial-lateral preoptic glutamatergic neurons altered EEG features during wakefulness and NREM sleep

This study aimed to determine if the activation of preoptic glutamatergic neurons, in addition to altering sleep-wake patterns, alters cortical dynamics (i.e., oscillations, connectivity, and complexity) associated with sleep and wakefulness. We thus evaluated local and network-level cortical activity by means of power spectrum, functional and directed connectivity between frontal and occipital cortices, as well as the complexity of EEG signals. Figure 11 shows the mean spectral power from frontal and occipital cortices during wakefulness and NREM sleep. There was a significant treatment effect (CNO) on the spectral power in the occipital cortex during wakefulness (F(1,9) = 7.38; p = 0.0238), and in the frontal and occipital cortices during NREM sleep [F(1,9) = 5.74; p = 0.0402 and F(1,9) = 7.26; p = 0.0246, respectively]. There was a significant drug by EEG frequency band interaction for the occipital cortex during wakefulness [F(6,54) = 8.89; p < 0.0001], as well as for frontal and occipital cortices during NREM sleep [F(6,54) = 7.92; p < 0.0001 and F(6,54) = 7.82; p < 0.0001, respectively]. No significant treatment effect or treatment by EEG frequency band interaction was found in the frontal cortex during wakefulness [F(1,9) = 1.91; p = 0.2002 and F(6,54) = 1.91; p = 0.0951]; there was a significant EEG frequency effect [F(6,54) = 61.3; p < 0.0001]. Post hoc multiple comparisons analysis revealed that CNO injection significantly decreased the power of slow oscillations (0.5 to 1 Hz) during wakefulness (p = 0.0041, frontal, and p < 0.0001, occipital) and NREM sleep (p < 0.0001, frontal, and p < 0.0001, occipital).

**Figure 11.**
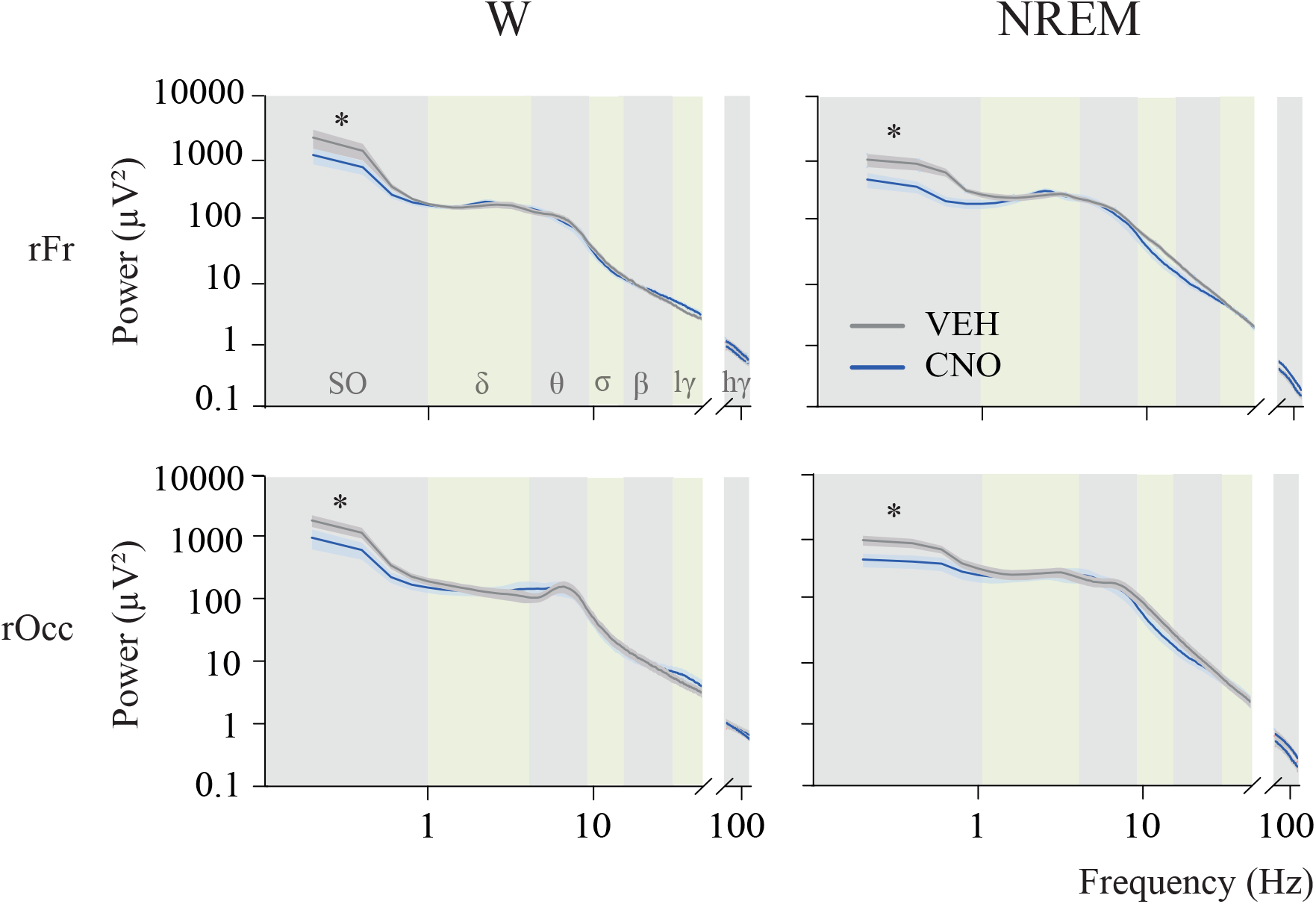
Activation of glutamatergic neurons in the medial-lateral preoptic region decreased the spectral power of slow oscillations. The graphs plot spectral power changes in the right frontal (rFr) and occipital (rOcc) regions during wakefulness (W) and NREM sleep. Traces represent mean values (thin, dark lines) ± the standard error of the mean (shaded area above and below the mean). Frequency ranges are indicated by alternating horizontal colored bands in the background of the graphs. Because notch filters were applied to some recording pairs (i.e., vehicle and CNO recordings from the same mouse), frequencies between 45 and 75 Hz were excluded from the analysis. Two-way repeated measures ANOVA followed by a Sidak test corrected for multiple comparisons were employed for statistical comparison of spectral power in each frequency band. Asterisks indicate significant differences (p < 0.05) relative to control. Abbreviations: SO, slow oscillations; δ, delta; θ, theta; σ, sigma; β, beta; lγ, low-gamma; hγ, high-gamma, VEH, vehicle and CNO, clozapine-N-oxide.

Figure 12 plots Z’coherence between frontal and occipital cortices as a function of EEG frequency band and treatment condition. ANOVA revealed a significant treatment by EEG frequency band interaction during wakefulness [F(6,54) = 3.34; p = 0.0072] and NREM sleep [F(6,54) = 6.13; p < 0.0001]. Furthermore, a post hoc multiple comparisons test demonstrated an increase in high-gamma coherence during wakefulness (p = 0.0449), as well as delta (p = 0.0163), theta (p < 0.0001), and high-gamma (p = 0.0209) Z’coherence during NREM sleep. In accordance with the changes in Z’coherence (undirected functional connectivity), CNO administration altered NSTE (directed connectivity). Mean feedforward and feedback NTSE values for each frequency band during wake and NREM states are shown in Table 2. In comparison to vehicle, CNO injection significantly altered feedforward connectivity during NREM sleep [F(1,9) = 6.60; p = 0.0302] but not during wakefulness [F(1,9) = 1.86; p = 0.2059]. Moreover, there was a significant drug by frequency band interaction for the frontal-to-occipital NSTE during NREM [F(5,45) = 4.05; p = 0.0040] and for the occipital-to-frontal NSTE during wakefulness [F(5,45) = 3.71; p = 0.0068] and NREM sleep [F(5,45) = 4.95; p = 0.0011]. Post hoc multiple comparisons tests demonstrated that theta frontal-to-occipital connectivity was increased during NREM sleep (p = 0.0003). Additionally, theta occipital-to-frontal connectivity was increased during NREM (p < 0.0001) and wakefulness (p = 0.0052). Compared to vehicle injection, CNO did not modify feedforward NSTE during NREM in delta (p = 0.0759), sigma, beta, low-gamma, and high-gamma (p > 0.9999 for each frequency band), as well as the feedback NSTE during wakefulness or NREM in delta (p = 0.8974 and p > 0.9999, respectively), sigma, beta, low-gamma, and high-gamma (p > 0.9999 for each frequency band and each behavioral state). To summarize, activation of preoptic glutamatergic neurons enhanced both undirected and directed connectivity in the theta frequency band during NREM sleep as well as undirected functional connectivity in the high-gamma band during wakefulness and NREM sleep.

**Figure 12.**
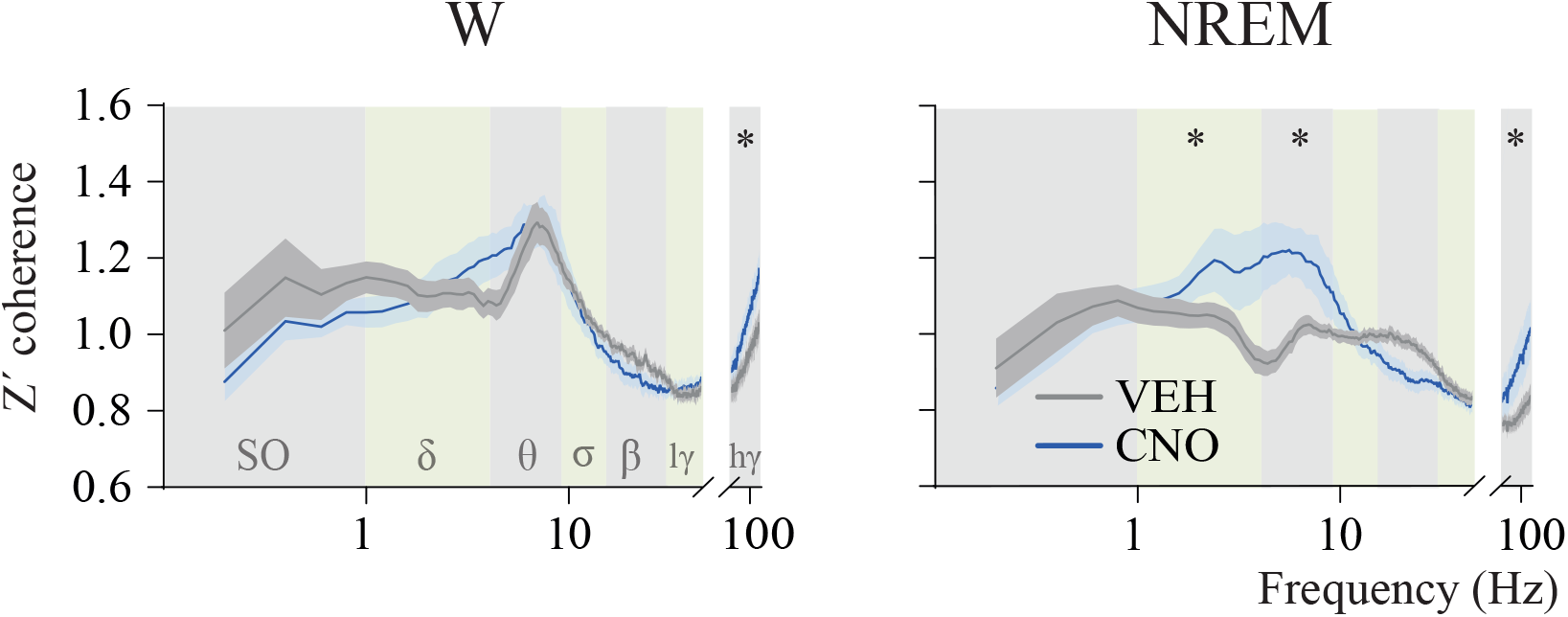
Activation of glutamatergic neurons in the medial-lateral preoptic region increased cortical connectivity. Mean z’coherence (undirected connectivity) profile between right frontal and occipital as a function of state (wakefulness (W) and NREM sleep) and treatment (vehicle (VEH) and clozapine-N-oxide (CNO)). Traces represent mean values (thin, dark lines) ± the standard error of the mean (shaded area above and below the mean). Frequency ranges are indicated by alternating horizontal colored bands in the background of the graphs. Because notch filters were applied to some recording pairs (i.e., vehicle and CNO recordings from the same mouse), frequencies between 45 and 75 Hz were excluded from the analysis. Asterisks indicate significant differences (p < 0.05) relative to control. Abbreviations: SO, slow oscillations; δ, delta; θ, theta; σ, sigma; β, beta; lγ, low-gamma; hγ, high-gamma, VEH, vehicle and CNO, clozapine-N-oxide.

**Table 2.** Directed cortical connectivity during NREM and wake states, by EEG frequency band and as a function of treatment condition.

|  | Occipital-to-Frontal<br>(Feedback Connectivity) |  |  |  | Frontal-to-Occipital<br>(Feedforward Connectivity) |  |  |  |
| --- | --- | --- | --- | --- | --- | --- | --- | --- |
|  | Wakefulness |  | NREM sleep |  | Wakefulness |  | NREM sleep |  |
| EEG Frequency band | VEH | CNO | VEH | CNO | VEH | CNO | VEH | CNO |
| Delta<br>(0.5 - 4 Hz) | 0.033<br>± 0.002 | 0.029<br>± 0.002 | 0.028<br>± 0.001 | 0.031<br>± 0.003 | 0.037<br>± 0.002 | 0.048<br>± 0.007 | 0.052<br>± 0.008 | 0.038<br>± 0.002 |
| Theta<br>(4 - 9 Hz) | 0.049<br>± 0.003 | 0.059 ±<br>0.004 | 0.035<br>± 0.001 | **0.051\***<br>± **0.004** | 0.086<br>± 0.006 | **0.090\***<br>± **0.009** | 0.073<br>± 0.010 | **0.050\***<br>± **0.003** |
| Sigma<br>(9 - 15 Hz) | 0.049<br>± 0.002 | 0.047<br>± 0.005 | 0.049<br>± 0.003 | 0.047<br>± 0.005 | 0.049<br>± 0.003 | 0.050<br>± 0.007 | 0.040<br>± 0.004 | 0.039<br>± 0.001 |
| Beta<br>(15 - 30 Hz) | 0.026<br>± 0.024 | 0.023<br>± 0.002 | 0.027<br>± 0.001 | 0.023<br>± 0.001 | 0.0250 ±<br>0.001 | 0.022<br>± 0.001 | 0.021<br>± 0.001 | 0.027<br>± 0.001 |
| LG<br>(30 - 45 Hz) | 0.021<br>± 0.001 | 0.024<br>± 0.002 | 0.022<br>± 0.001 | 0.023<br>± 0.001 | 0.018<br>± 0.001 | 0.019<br>± 0.001 | 0.019<br>± 0.001 | 0.019<br>± 0.001 |
| HG<br>(75 - 115 Hz) | 0.017<br>± 0.001 | 0.020<br>± 0.001 | 0.011<br>± 0.001 | 0.015<br>± 0.002 | 0.020<br>± 0.001 | 0.025<br>± 0.001 | 0.019<br>± 0.002 | 0.014<br>± 0.001 |

Data are presented as mean ± standard error of the mean. A two-way repeated measures ANOVA followed by Sidak multiple comparisons test was used for statistical comparisons. Asterisks and bold fonts indicate significant differences relative to control ( $p < 0.05$ ). CNO - clozapine-N-oxide (agonist at hM3Dq receptors), HG - high gamma, LG - low gamma, VEH - vehicle.

Last, changes in temporal complexity of the signals evaluated by a corrected Lempel-Ziv Complexity analysis are summarized in Figure 13. Relative to vehicle injection, CNO increased EEG signal complexity during wakefulness in both frontal [Mean ± SEM = 0.9782 ± 0.005 vs 0.9618 ± 0.005; t(9) = 2.47, p = 0.0354] and occipital regions [Mean ± SEM = 0.9891 ± 0.007 vs 0.968 ± 0.007; t(9) = 3.72, p = 0.0048]. Additionally, CNO increased the EEG complexity during NREM sleep in frontal [Mean ± SEM = 0.9728 ± 0.009 vs 0.947 ± 0.003; t(9) = 3.29, p = 0.0094], and occipital regions [Mean ± SEM = 0.9708 ± 0.009 vs 0.948 ± 0.003; t(9) = 3.13, p = 0.0122]. Interestingly, EEG complexity levels during NREM sleep after CNO were not significantly different than during wakefulness in the control treatment condition (frontal cortex: t(9) = 0.94; p = 0.3719 and occipital cortex: t(9) = 0.26 p = 0.8029).

**Figure 13.**
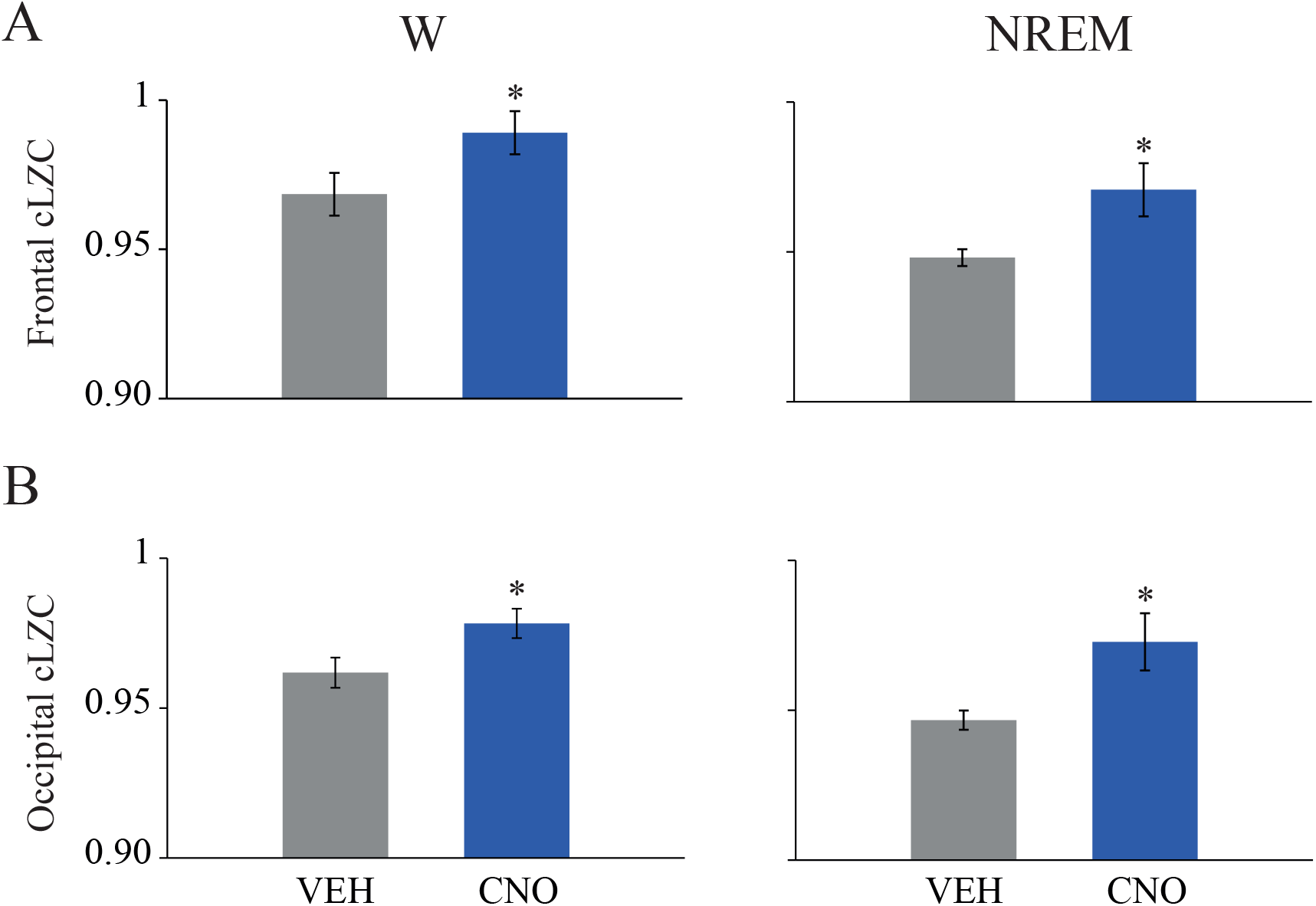
Activation of glutamatergic neurons in the medial-lateral preoptic region increased electroencephalographic (EEG) signal complexity. The graphs plot corrected Lempel-Ziv complexity (cLZC) values during wakefulness (W) and NREM sleep for frontal (*A*) and occipital (*B*) regions. Data are reported as the mean ± standard error of the mean. Differences between vehicle (VEH) and clozapine-N-oxide (CNO) were analyzed by two-tailed paired t-tests. Asterisks indicate significant differences (p < 0.05) relative to control.

## Discussion

This study demonstrates that chemogenetic activation of preoptic glutamatergic neurons within the medial-lateral preoptic region of the hypothalamus increases wakefulness, decreases NREM sleep, causes NREM sleep fragmentation and state instability, and suppresses REM sleep. Furthermore, these neurons influence cortical oscillations, connectivity, and complexity. A dual role for preoptic neurons in sleep-wake regulation was recently predicted by modeling simulations based on EEG dynamics across the sleep-wake cycle of rats with VLPO neuronal lesions (Lombardi et al., 2020). Together with our previous work (Vanini et al., 2020), and evidence demonstrating that optogenetic stimulation of VLPO glutamatergic neurons projecting to the tuberomammillary nucleus increases wakefulness (Chung et al., 2017), this is among the first studies to empirically identify a subset of preoptic neurons that promotes wakefulness. Importantly, activation of glutamatergic neurons in the MnPO (used here as a control site) had no substantial effect on sleep-wake duration, sleep latency, and consolidation. The non-significant trend towards an increase in the latency and decrease in the duration of REM sleep during the first 4 hours post-CNO injection is congruent with previous work from our group showing that (over 3 hours) MnPO glutamatergic neurons only reduced REM sleep quantity (Vanini et al., 2020). Together, these data suggest that glutamatergic neurons in the MnPO destabilize REM sleep and support the conclusion that the effect of the activation of glutamatergic neurons in the medial-lateral preoptic region on sleep-wake states is site-specific.

The preoptic area contributes to REM sleep regulation. Cell-specific excitotoxic lesions in a region medial and dorsal to the VLPO (termed “extended” VLPO) decreases REM sleep (Lu et al., 2002). Subsets of preoptic neurons are maximally active during spontaneous REM sleep bouts, during periods of sleep restriction when REM sleep pressure is the highest, and during REM sleep rebound following REM sleep deprivation (Lu et al., 2002; Suntsova et al., 2002; Gvilia et al., 2006; Takahashi et al., 2009; Sakai, 2011; Alam et al., 2014). Furthermore, activation of preoptic GABAergic neurons that innervate the tuberomammillary nucleus increases REM sleep (Chung et al., 2017). Here we show that activation of glutamatergic neurons in the medial-lateral preoptic region decreased REM sleep time. In fact, this state was eliminated for six hours in 55% of the mice. To the best of our knowledge, this is the first direct demonstration that preoptic glutamatergic neurons can gate REM sleep generation. The remarkable reduction in REM sleep may result from several factors. First, the profound sleep fragmentation induced by activation of preoptic glutamatergic neurons can reduce REM sleep because its propensity builds up during prior NREM sleep bouts and spontaneous REM sleep generation requires NREM sleep occurrence (Benington and Heller, 1994; Le Bon, 2020). However, while NREM sleep fragmentation had a duration of four hours, REM sleep quantity was reduced or totally suppressed for six hours post-CNO administration and there was no negative correlation between REM sleep time and sleep fragmentation or the number of NREM-wake transitions. Furthermore, there was no REM homeostatic response after sleep fragmentation, as both number and duration of REM sleep bouts remained unchanged. Another potential mechanism is by a direct or indirect activation of brain circuits that inhibit REM sleep generation. The preoptic region is reciprocally connected with wake-promoting monoaminergic systems and hypocretin/orexin-containing neurons in the perifornical region of the hypothalamus (Zardetto-Smith and Johnson, 1995; Sherin et al., 1998; Chou et al., 2002; Uschakov et al., 2006). Furthermore, the ventrolateral periaqueductal gray receives projections from preoptic neurons (Uschakov et al., 2009; Hsieh et al., 2011) and GABAergic neurons within the periaqueductal gray region exert a powerful inhibitory control on REM sleep generation (Sastre et al., 1996; Vanini et al., 2007; Sapin et al., 2009; Weber et al., 2018). Identification of local and distant target neurons of wake-promoting glutamatergic neurons in the medial-lateral preoptic region will be critical for a circuit-based understanding of their role in REM sleep control. This is especially important because the ability of these neurons to promote wakefulness as well as enhance cortical dynamics while suppressing REM sleep might serve the function of preventing REM-like intrusions after the initiation of waking consciousness. Last, preoptic neurons play an important role in thermoregulation (McGinty et al., 2001). Activation of MnPO glutamatergic (Abbott and Saper, 2017; Vanini et al., 2020) and VLPO galaninergic neurons (Kroeger et al., 2018) induces significant hypothermia in mice, and exposure to cold temperature increases wakefulness and reduces REM sleep in rodents (Roussel et al., 1984; Amici et al., 1998; Baracchi et al., 2008; Sato et al., 2015). Glutamatergic neurons that cause hypothermia (some of which also promote sleep) are mainly localized within the MnPO and medial preoptic area (Zhang et al., 2015; Abbott and Saper, 2017; Harding et al., 2018; Vanini et al., 2020). By contrast, activation of glutamatergic neurons in the medial-lateral preoptic region suppressed REM sleep but did not alter core body temperature.

This study shows that activation of glutamatergic neurons in the medial-lateral preoptic region causes a transient increase in wakefulness and a reduction in NREM sleep. Additionally, there was a robust, long-lasting NREM sleep fragmentation and state instability resembling the disrupted sleep pattern observed in sleep apnea (Guilleminault et al., 1976; Kimoff, 1996), Alzheimer’s disease (Lim et al., 2014), and aging (Lim et al., 2014; Li et al., 2018). Similarly, chemogenetic stimulation of cholinergic neurons in the basal forebrain, lateral to the preoptic area, increases wakefulness and fragments sleep (Anaclet et al., 2015) and optical stimulation of basal forebrain parvalbumin neurons induces brief, rapid arousals from sleep (McKenna et al., 2020). Relative to basal forebrain glutamatergic neurons, optogenetic stimulation increases wakefulness (Xu et al., 2015), whereas activation of these neurons using chemogenetic strategies did not significantly alter sleep-wake states (Anaclet et al., 2015). Importantly, none of the mice in this study expressed excitatory designer receptors in the basal forebrain region, confirming again the specificity of the effects of our stimulation region. Given the frequent and brief arousals from sleep observed after CNO administration, we speculate that glutamatergic neurons in the medial-lateral preoptic region may initiate but not maintain wakefulness. Furthermore, these neurons may be part of a brain network that produces rapid arousals from sleep in response to endogenous or environmental alert signals.

Glutamatergic neurons in the medial-lateral preoptic altered several sleep and wake EEG features related to cortical dynamics. Specifically, CNO administration increased (1) theta and gamma coherence, (2) cortical connectivity, and (3) EEG complexity during NREM sleep. These are electrophysiologic traits of more activated states. Indeed, EEG theta and gamma coherence, directed connectivity and complexity are typically highest during wakefulness (Abasolo et al., 2015; Pal et al., 2016; Brito et al., 2020; Mondino et al., 2020; Pal et al., 2020) and progressively decline as sleep deepens (Abasolo et al., 2015; Pal et al., 2016; Bandt, 2017; Gonzalez et al., 2019; Migliorelli et al., 2019; Gonzalez et al., 2020). Together, these EEG changes and the reduction of the spectral power of slow oscillations after CNO injection provide evidence for a “lighter” NREM sleep state. Increased EEG complexity and high-gamma coherence are, respectively, suggestive of a higher number of brain network interactions (Guevara Erra et al., 2016) and high alertness during wakefulness (Castro et al., 2013). In the current study, we indeed found that CNO administration increased high-gamma coherence, EEG complexity, and feedback connectivity, while reducing the power of slow oscillations during wakefulness. Therefore, these data support the interpretation that activation of glutamatergic neurons in the lateral-medial preoptic region produces an activated EEG state with features that correlate with enhanced wakefulness. Sleep consolidation (Ward et al., 2009) and sleep quality (i.e., slow wave activity and slow oscillations during NREM sleep) (Bellesi et al., 2014; Schreiner et al., 2018; Kim et al., 2019; Hahn et al., 2020), as well as REM sleep (Boyce et al., 2017) are crucial for normal cognition. Thus, because of the substantial disruption of sleep patterns and cortical dynamics produced during NREM sleep by the activation of preoptic glutamatergic neurons, further studies are needed to examine the impact on attention and memory processes.

Collectively, our results support the interpretation that glutamatergic neurons in the medial-lateral preoptic region may initiate (but not maintain) wakefulness from sleep and their inactivation may be necessary for NREM stability and REM sleep initiation. The present work is limited by including only male mice and stimulation strategies as well as by the lack of data on the effects of these preoptic neurons on sleep and wake states during the dark phase. Additionally, future studies are needed to identify relevant projection pathways and neuronal targets mediating their wakefulness-promoting effects. Furthermore, these findings encourage future circuit-based studies to examine the relevance of these neurons in the generation of wakefulness in response to endogenous or environmental alert signals during sleep, while suppressing REM-like intrusions. These data also have translational implications as they potentially inform the etiology of debilitating, disrupted sleep patterns observed in aging, sleep apnea, and dementia. Last, this study advances sleep neurobiology by providing empirical evidence supporting the notion that the preoptic area is not exclusively somnogenic but rather plays a dual role in the regulation of both sleep and wakefulness.

## Acknowledgments

The authors thank Leonor Floran-Garduno, Megumi Mast and Mary Norat from the Department of Anesthesiology, University of Michigan for expert assistance. This work was funded by the National Institutes of Health (R01 GM124248 to G.V. and G.A.M.), Bethesda, MD, USA, and the Department of Anesthesiology, University of Michigan Medical School, Ann Arbor.

## References

Abasolo D, Simons S, Morgado da Silva R, Tononi G, Vyazovskiy VV (2015) Lempel-Ziv complexity of cortical activity during sleep and waking in rats. J Neurophysiol 113:2742–2752.

Abbott SBG, Saper CB (2017) Median preoptic glutamatergic neurons promote thermoregulatory heat loss and water consumption in mice. J Physiol 595:6569–6583.

Alam MA, Kumar S, McGinty D, Alam MN, Szymusiak R (2014) Neuronal activity in the preoptic hypothalamus during sleep deprivation and recovery sleep. J Neurophysiol 111:287–299.

Alam MN, Mallick BN (1990) Differential acute influence of medial and lateral preoptic areas on sleep-wakefulness in freely moving rats. Brain Res 525:242–248.

Amici R, Zamboni G, Perez E, Jones CA, Parmeggiani PL (1998) The influence of a heavy thermal load on REM sleep in the rat. Brain Res 781:252–258.

Anaclet C, Pedersen NP, Ferrari LL, Venner A, Bass CE, Arrigoni E, Fuller PM (2015) Basal forebrain control of wakefulness and cortical rhythms. Nat Commun 6:8744.

Bandt C (2017) A New Kind of Permutation Entropy Used to Classify Sleep Stages from Invisible EEG Microstructure. Entropy 19.

Baracchi F, Zamboni G, Cerri M, Del Sindaco E, Dentico D, Jones CA, Luppi M, Perez E, Amici R (2008) Cold exposure impairs dark-pulse capacity to induce REM sleep in the albino rat. J Sleep Res 17:166–179.

Bellesi M, Riedner BA, Garcia-Molina GN, Cirelli C, Tononi G (2014) Enhancement of sleep slow waves: underlying mechanisms and practical consequences. Front Syst Neurosci 8:208.

Benedetto L, Chase MH, Torterolo P (2012) GABAergic processes within the median preoptic nucleus promote NREM sleep. Behav Brain Res 232:60–65.

Benington J, Heller HC (1994) REM-sleep timing is controlled homeostatically by accumulation of REM-sleep propensity in non-REM sleep. Am J Physiol Regul Integr Comp Physiol 266:1992–2000.

Borjigin J, Lee U, Liu T, Pal D, Huff S, Klarr D, Sloboda J, Hernandez J, Wang MM, Mashour GA (2013) Surge of neurophysiological coherence and connectivity in the dying brain. Proc Natl Acad Sci U S A 110:14432–14437.

Boyce R, Williams S, Adamantidis A (2017) REM sleep and memory. Curr Opin Neurobiol 44:167–177.

Brito MA, Li D, Mashour GA, Pal D (2020) State-Dependent and Bandwidth-Specific Effects of Ketamine and Propofol on Electroencephalographic Complexity in Rats. Front Syst Neurosci 14:50.

Castro S, Falconi A, Chase MH, Torterolo P (2013) Coherent neocortical 40-Hz oscillations are not present during REM sleep. Eur J Neurosci 37:1330–1339.

Chou TC, Bjorkum AA, Gaus SE, Lu J, Scammell TE, Saper CB (2002) Afferents to the ventrolateral preoptic nucleus. J Neurosci 22:977–990.

Chung S, Weber F, Zhong P, Tan CL, Nguyen TN, Beier KT, Hormann N, Chang WC, Zhang Z, Do JP, Yao S, Krashes MJ, Tasic B, Cetin A, Zeng H, Knight ZA, Luo L, Dan Y (2017) Identification of preoptic sleep neurons using retrograde labelling and gene profiling. Nature 545:477–481.

Delorme A, Makeig S (2004) EEGLAB: an open source toolbox for analysis of single-trial EEG dynamics including independent component analysis. J Neurosci Methods 134:9–21.

Dentico D, Amici R, Baracchi F, Cerri M, Del Sindaco E, Luppi M, Martelli D, Perez E, Zamboni G (2009) c-Fos expression in preoptic nuclei as a marker of sleep rebound in the rat. Eur J Neurosci 30:651–661.

Eikermann M, Vetrivelan R, Grosse-Sundrup M, Henry ME, Hoffmann U, Yokota S, Saper CB, Chamberlin NL (2011) The ventrolateral preoptic nucleus is not required for isoflurane general anesthesia. Brain Res 1426:30–37.

Gaus SE, Strecker RE, Tate BA, Parker RA, Saper CB (2002) Ventrolateral preoptic nucleus contains sleep-active, galaninergic neurons in multiple mammalian species. Neuroscience 115:285–294.

Gonzalez J, Cavelli M, Mondino A, Pascovich C, Castro-Zaballa S, Torterolo P, Rubido N (2019) Decreased electrocortical temporal complexity distinguishes sleep from wakefulness. Sci Rep 9:18457.

Gonzalez J, Cavelli M, Mondino A, Pascovich C, Castro-Zaballa S, Rubido N, Torterolo P (2020) Electrocortical temporal complexity during wakefulness and sleep: an updated account. In: Sleep Science, pp 47–50.

Guevara Erra R, Mateos DM, Wennberg R, Perez Velazquez JL (2016) Towards a statistical mechanics of consciousness: maximization of number of connections is associated with conscious awareness. Physical Review E 94.

Guilleminault C, Tilkian A, Dement WC (1976) The sleep apnea syndromes. Annu Rev Med 27:465–484.

Gvilia I, Turner A, McGinty D, Szymusiak R (2006) Preoptic area neurons and the homeostatic regulation of rapid eye movement sleep. J Neurosci 26:3037–3044.

Gvilia I, Suntsova N, Kostin A, Kalinchuk A, McGinty D, Basheer R, Szymusiak R (2017) The role of adenosine in the maturation of sleep homeostasis in rats. J Neurophysiol 117:327–335.

Hahn MA, Heib D, Schabus M, Hoedlmoser K, Helfrich RF (2020) Slow oscillation-spindle coupling predicts enhanced memory formation from childhood to adolescence. Elife 9.

Harding EC, Yu X, Miao A, Andrews N, Ma Y, Ye Z, Lignos L, Miracca G, Ba W, Yustos R, Vyssotski AL, Wisden W, Franks NP (2018) A Neuronal Hub Binding Sleep Initiation and Body Cooling in Response to a Warm External Stimulus. Curr Biol 28:2263–2273 e2264.

Hsieh KC, Gvilia I, Kumar S, Uschakov A, McGinty D, Alam MN, Szymusiak R (2011) c-Fos expression in neurons projecting from the preoptic and lateral hypothalamic areas to the ventrolateral periaqueductal gray in relation to sleep states. Neuroscience 188:55–67.

Ilg AK, Enkel T, Bartsch D, Bahner F (2018) Behavioral Effects of Acute Systemic Low-Dose Clozapine in Wild-Type Rats: Implications for the Use of DREADDs in Behavioral Neuroscience. Front Behav Neurosci 12:173.

John J, Kumar VM, Gopinath G (1998) Recovery of sleep after fetal preoptic transplantation in medial preoptic area-lesioned rats. Sleep 21:601–606.

John J, Kumar VM, Gopinath G, Ramesh V, Mallick H (1994) Changes in sleep-wakefulness after kainic acid lesion of the preoptic area in rats. Jpn J Physiol 44:231–242.

Jordan D, Ilg R, Riedl V, Schorer A, Grimberg S, Neufang S, Omerovic A, Berger S, Untergehrer G, Preibisch C, Schulz E, Schuster T, Schroter M, Spoormaker V, Zimmer C, Hemmer B, Wohlschlager A, Kochs EF, Schneider G (2013) Simultaneous electroencephalographic and functional magnetic resonance imaging indicate impaired cortical top-down processing in association with anesthetic-induced unconsciousness. Anesthesiology 119:1031–1042.

Kim J, Gulati T, Ganguly K (2019) Competing Roles of Slow Oscillations and Delta Waves in Memory Consolidation versus Forgetting. Cell 179:514–526 e513.

Kim JW, Lee JS, Robinson PA, Jeong DU (2009) Markov analysis of sleep dynamics. Phys Rev Lett 102:178104.

Kimoff J (1996) Sleep Fragmentation In Obstructive Sleep Apnea. Sleep 19:61–66.

Koyama Y, Hayaishi O (1994) Firing of neurons in the preoptic/anterior hypothalamic areas in rat: its possible involvement in slow wave sleep and paradoxical sleep. Neurosci Res 19:31–38.

Krashes MJ, Koda S, Ye C, Rogan SC, Adams AC, Cusher DS, Maratos-Flier E, Roth BL, Lowell BB (2011) Rapid, reversible activation of AgRP neurons drives feeding behavior in mice. J Clin Invest 121:1424–1428.

Kroeger D, Absi G, Gagliardi C, Bandaru SS, Madara JC, Ferrari LL, Arrigoni E, Munzberg H, Scammell TE, Saper CB, Vetrivelan R (2018) Galanin neurons in the ventrolateral preoptic area promote sleep and heat loss in mice. Nat Commun 9:4129.

Le Bon O (2020) Relationships between REM and NREM in the NREM-REM sleep cycle: a review on competing concepts. Sleep Med 70:6–16.

Lee U, Ku S, Noh G, Baek S, Choi B, Mashour GA (2013) Disruption of frontal-parietal communication by ketamine, propofol, and sevoflurane. Anesthesiology 118:1264–1275.

Lempel A, Ziv J (1976) On the complexity of finite sequences. IEEE Trans Inf Theory:75–81.

Li D, Mashour GA (2019) Cortical dynamics during psychedelic and anesthetized states induced by ketamine. Neuroimage 196:32–40.

Li D, Hambrecht-Wiedbusch VS, Mashour GA (2017) Accelerated Recovery of Consciousness after General Anesthesia Is Associated with Increased Functional Brain Connectivity in the High-Gamma Bandwidth. Front Syst Neurosci 11:16.

Li J, Vitiello MV, Gooneratne NS (2018) Sleep in Normal Aging. Sleep Med Clin 13:1–11.

Lim AS, Ellison BA, Wang JL, Yu L, Schneider JA, Buchman AS, Bennett DA, Saper CB (2014) Sleep is related to neuron numbers in the ventrolateral preoptic/intermediate nucleus in older adults with and without Alzheimer's disease. Brain 137:2847–2861.

Lin JS, Sakai K, Vanni-Mercier G, Jouvet M (1989) A critical role of the posterior hypothalamus in the mechanisms of wakefulness determined by microinjection of muscimol in freely moving cats. Brain Res 479:225–240.

Lombardi F, Gomez-Extremera M, Bernaola-Galvan P, Vetrivelan R, Saper CB, Scammell TE, Ivanov PC (2020) Critical Dynamics and Coupling in Bursts of Cortical Rhythms Indicate Non-Homeostatic Mechanism for Sleep-Stage Transitions and Dual Role of VLPO Neurons in Both Sleep and Wake. J Neurosci 40:171–190.

Lu J, Greco MA, Shiromani P, Saper CB (2000) Effect of lesions of the ventrolateral preoptic nucleus on NREM and REM sleep. J Neurosci 20:3830–3842.

Lu J, Bjorkum AA, Xu M, Gaus SE, Shiromani PJ, Saper CB (2002) Selective activation of the extended ventrolateral preoptic nucleus during rapid eye movement sleep. J Neurosci 22:4568–4576.

Ma Y, Miracca G, Yu X, Harding EC, Miao A, Yustos R, Vyssotski AL, Franks NP, Wisden W (2019) Galanin Neurons Unite Sleep Homeostasis and alpha2-Adrenergic Sedation. Curr Biol 29:3315–3322 e3313.

McGinty D, Alam MN, Szymusiak RN M., Yamamoto M (2001) Hypothalamic Sleep-Promoting Mechanisms: Coupling to thermorregulation. Archives Italiannes de Biologie 139:63–75.

McKenna JT, Thankachan S, Uygun DS, Shukla C, McNally JM, Schiffino FL, Cordeira J, Katsuki F, Zant JC, Gamble MC, Deisseroth K, McCarley RW, Brown RE, Strecker RE, Basheer R (2020) Basal Forebrain Parvalbumin Neurons Mediate Arousals from Sleep Induced by Hypercarbia or Auditory Stimuli. Curr Biol 30:2379–2385 e2374.

Mendelson WB (2000) Sleep-inducing effects of adenosine microinjections into the medial preoptic area are blocked by flumazenil. Brain Res 852:479–481.

Migliorelli C, Bachiller A, Andrade AG, Alonso JF, Mananas MA, Borja C, Gimenez S, Antonijoan RM, Varga AW, Osorio RS, Romero S (2019) Alterations in EEG connectivity in healthy young adults provide an indicator of sleep depth. Sleep 42.

Miranda de Sá AMFL, Ferreira DD, Dias E, Mendes EMAM, Felix LB (2009) Coherence estimate between a random and a periodic signal: Bias, variance, analytical critical values, and normalizing transforms. Journal of the Franklin Institute 346:841–853.

Mitra P, Bokil H (2008) Observed Brain Dynamics. New York: Oxford University Press.

Mondino A, Cavelli M, Gonzalez J, Osorio L, Castro-Zaballa S, Costa A, Vanini G, Torterolo P (2020) Power and coherence in the EEG of the rat: impact of behavioral states, cortical area, lateralization and light/dark phases. bioRxiv preprint.

Nauta WJ (1946) Hypothalamic regulation of sleep in rats; an experimental study. J Neurophysiol 9:285–316.

Pal D, Silverstein BH, Lee H, Mashour GA (2016) Neural Correlates of Wakefulness, Sleep, and General Anesthesia: An Experimental Study in Rat. Anesthesiology 125:929–942.

Pal D, Li D, Dean JG, Brito MA, Liu T, Fryzel AM, Hudetz AG, Mashour GA (2020) Level of Consciousness Is Dissociable from Electroencephalographic Measures of Cortical Connectivity, Slow Oscillations, and Complexity. J Neurosci 40:605–618.

Parmeggiani PL (1987) Interaction between sleep and thermoregulation: An aspect of the Control of Behavioral States. Sleep 10:426–435.

Paxinos G K.B.J.F (2001) The Mouse Brain in Stereotaxic Coordinates, 2 Edition. San Diego: Academic.

Perez-Atencio L, Garcia-Aracil N, Fernandez E, Barrio LC, Barios JA (2018) A four-state Markov model of sleep-wakefulness dynamics along light/dark cycle in mice. PLoS One 13:e0189931.

Ranft A, Golkowski D, Kiel T, Riedl V, Kohl P, Rohrer G, Pientka J, Berger S, Thul A, Maurer M, Preibisch C, Zimmer C, Mashour GA, Kochs EF, Jordan D, Ilg R (2016) Neural Correlates of Sevoflurane-induced Unconsciousness Identified by Simultaneous Functional Magnetic Resonance Imaging and Electroencephalography. Anesthesiology 125:861–872.

Rezai Amin S, Gruszczynski C, Guiard BP, Callebert J, Launay JM, Louis F, Betancur C, Vialou V, Gautron S (2019) Viral vector-mediated Cre recombinase expression in substantia nigra induces lesions of the nigrostriatal pathway associated with perturbations of dopamine-related behaviors and hallmarks of programmed cell death. J Neurochem 150:330–340.

Roussel B, Turrillot P, Kitahama K (1984) Effect of ambient temperature on the sleep-waking cycle in two strains of mice. Brain Res 294:67–73.

Sakai K (2011) Sleep-waking discharge profiles of median preoptic and surrounding neurons in mice. Neuroscience 182:144–161.

Sallanon M, Denoyer M, Kitahama K, Aubert C, Gay N, Jouvet M (1989) Long-lasting insomnia induced by preoptic neuron lesions and its transient reversal by muscimol injection into the posterior hypothalamus in the cat. Neuroscience 32:669–683.

Sapin E, Lapray D, Berod A, Goutagny R, Leger L, Ravassard P, Clement O, Hanriot L, Fort P, Luppi PH (2009) Localization of the brainstem GABAergic neurons controlling paradoxical (REM) sleep. PLoS One 4:e4272.

Sastre JP, Buda C, Kitahama K, Jouvet M (1996) Importance of the ventrolateral region of the periaqueductal gray and adjacent tegmentum in the control of paradoxical sleep as studied by muscimol microinjections in the cat. Neuroscience 74:415–426.

Sato N, Marui S, Ozaki M, Nagashima K (2015) Cold exposure and/or fasting modulate the relationship between sleep and body temperature rhythms in mice. Physiol Behav 149:69–75.

Schartner M, Seth A, Noirhomme Q, Boly M, Bruno MA, Laureys S, Barrett A (2015) Complexity of Multi-Dimensional Spontaneous EEG Decreases during Propofol Induced General Anaesthesia. PLoS One 10:e0133532.

Schartner MM, Carhart-Harris RL, Barrett AB, Seth AK, Muthukumaraswamy SD (2017) Increased spontaneous MEG signal diversity for psychoactive doses of ketamine, LSD and psilocybin. Sci Rep 7:46421.

Schreiner T, Doeller CF, Jensen O, Rasch B, Staudigl T (2018) Theta Phase-Coordinated Memory Reactivation Reoccurs in a Slow-Oscillatory Rhythm during NREM Sleep. Cell Rep 25:296–301.

Sherin JE, Elmquist JK, Torrealba F, Saper CB (1998) Innervation of histaminergic tuberomammillary neurons by GABAergic and galaninergic neurons in the ventrolateral preoptic nucleus of the rat. J Neurosci 18:4705–4721.

Sitt JD, King JR, El Karoui I, Rohaut B, Faugeras F, Gramfort A, Cohen L, Sigman M, Dehaene S, Naccache L (2014) Large scale screening of neural signatures of consciousness in patients in a vegetative or minimally conscious state. Brain 137:2258–2270.

Suntsova N, Szymusiak R, Alam MN, Guzman-Marin R, McGinty D (2002) Sleep-waking discharge patterns of median preoptic nucleus neurons in rats. J Physiol 543:665–677.

Szymusiak R, Alam N, Steininger TL, McGinty D (1998) Sleep-waking discharge patterns of ventrolateral preoptic/anterior hypothalamic neurons in rats. Brain Res 803:178–188.

Takahashi K, Lin JS, Sakai K (2009) Characterization and mapping of sleep-waking specific neurons in the basal forebrain and preoptic hypothalamus in mice. Neuroscience 161:269–292.

Ticho SR, Radulovacki M (1991) Role of adenosine in sleep and temperature regulation in the preoptic area of rats. Pharmacol Biochem Behav 40:33–40.

Todd WD, Gibson JL, Shaw CS, Blumberg MS (2010) Brainstem and hypothalamic regulation of sleep pressure and rebound in newborn rats. Behav Neurosci 124:69–78.

Uschakov A, Gong H, McGinty D, Szymusiak R (2006) Sleep-active neurons in the preoptic area project to the hypothalamic paraventricular nucleus and perifornical lateral hypothalamus. Eur J Neurosci 23:3284–3296.

Uschakov A, McGinty D, Szymusiak R, McKinley MJ (2009) Functional correlates of activity in neurons projecting from the lamina terminalis to the ventrolateral periaqueductal gray. Eur J Neurosci 30:2347–2355.

Vanini G, Baghdoyan HA (2013) Extrasynaptic GABAA receptors in rat pontine reticular formation increase wakefulness. Sleep 36:337–343.

Vanini G, Torterolo P, McGregor R, Chase MH, Morales FR (2007) GABAergic processes in the mesencephalic tegmentum modulate the occurrence of active (rapid eye movement) sleep in guinea pigs. Neuroscience 145:1157–1167.

Vanini G, Bassana M, Mast M, Mondino A, Cerda I, Phyle M, Chen V, Colmenero AV, Hambrecht-Wiedbusch VS, Mashour GA (2020) Activation of Preoptic GABAergic or Glutamatergic Neurons Modulates Sleep-Wake Architecture, but Not Anesthetic State Transitions. Curr Biol 30:779–787 e774.

von Economo C (1930) Sleep as a problem of localization. The Journal of Nervous and Mental Disease:249–259.

Ward CP, McCoy JG, McKenna JT, Connolly NP, McCarley RW, Strecker RE (2009) Spatial learning and memory deficits following exposure to 24 h of sleep fragmentation or intermittent hypoxia in a rat model of obstructive sleep apnea. Brain Res 1294:128–137.

Weber F, Hoang Do JP, Chung S, Beier KT, Bikov M, Saffari Doost M, Dan Y (2018) Regulation of REM and Non-REM Sleep by Periaqueductal GABAergic Neurons. Nat Commun 9:354.

Xu M, Chung S, Zhang X, Zhong P, Ma C, Chang WC, Weissbourd B, Sakai N, Luo L, Nishino S, Dan Y (2015) Basal Forebrain Circuit for Sleep-Wake Control. Nat Neurosci 18.

Zardetto-Smith AM, Johnson AK (1995) Chemical topography of efferent projections from the median preoptic nucleus to pontine monoaminergic cell groups in the rat. Neurosci Lett 199:215–219.

Zhang Z, Ferretti V, Guntan I, Moro A, Steinberg EA, Ye Z, Zecharia AY, Yu X, Vyssotski AL, Brickley SG, Yustos R, Pillidge ZE, Harding EC, Wisden W, Franks NP (2015) Neuronal ensembles sufficient for recovery sleep and the sedative actions of alpha2 adrenergic agonists. Nat Neurosci 18:553–561.

